# Exploration of ligand (un)binding trajectories with Rosetta

**DOI:** 10.64898/2026.10.07.757317

**Authors:** Alexander Zlobin, Paul Eisenhuth, Valerio Ferrario, Jens Meiler

**Affiliations:** Institute for Drug Discovery, Leipzig University, 04103, Leipzig, Germany; Interdisciplinary Center for Bioinformatics, Leipzig University, 04107 Leipzig, Germany; Center for Scalable Data Analytics and Artificial Intelligence (ScaDS.AI) Dresden/Leipzig, Leipzig University, 04105, Leipzig, Germany; BASF SE, 67056 Ludwigshafen am Rhein, Germany; Center for Structural Biology, Vanderbilt University, Nashville, TN, 37235, USA; Department of Chemistry, Department of Pharmacology and Institute of Chemical Biology, Vanderbilt University, Nashville, TN, 37235, USA; School of Embedded Composite Artificial Intelligence SECAI, 04105, Leipzig, Germany; Faculty of Mathematics and Informatics, Faculty of Chemistry, Leipzig University, 04103, Leipzig, Germany

## Abstract

Ligand transport through proteins underlies many aspects of molecular function, but its detailed mechanisms remain largely inaccessible to direct observation. Molecular dynamics simulations could close this gap, but are computationally too demanding for practical purposes, including filtering stages of enzyme design pipelines. Simplified docking-based approaches might be informative, but sometimes the oversimplifications required for high throughput applications might lead to unreliable predictions. Here we present RosettaLigand PathFinder (RLPF), a fast and practical tool explicitly built to ensure conformational continuity of the ligand trajectory, protein and ligand flexibility, and robustness of every run. Built using a flexible RosettaScripts XML interface, it enables fine-grained control over all simulation parameters and adaptability for a wide range of use cases. Applied to several representative systems, RLPF reproduced established transport features and mutation effects, highlighting its ability to produce biologically relevant insights and serve as a filter in enzyme design campaigns. Wide customizability and well-defined scope of RLPF make it a useful addition to the family of tools to study the dynamics of interactions between small molecules and proteins.

## Introduction

Understanding the relationship between a protein sequence and its function is a central question of protein science. An established paradigm is to view biomolecules through the lens of physics, as mechanisms with structures and motions. This way, a function requires a particular structure, which is in turn encoded in a particular sequence. Such a relationship is crucial to uncover routes of diseases, develop drugs, find or engineer biocatalysts for pressing industrial needs ^1,2^, including the transition to green chemistry. Mapping enzyme structure-function relationships would allow us to understand effects of mutations, do structure-based rational design or condition high-throughput computational efforts ^3,4^, and create computational filters for such. To achieve that, it is important to study what structural features influence enzymatic activity and how. Well-established concepts here include active site preorganization, evolved electrostatic fields ^5,6^, selectivity through tighter binding of a cognate substrate or via programmed on-off switches, controlled tuning of enzymatic activity through allostery ^7^ and substrate or product inhibition ^8^, and more. A topic that is currently gaining momentum is the study of how an enzyme substrate or a product binds or unbinds from the active site, what features – including tunnels – are involved, and what associated challenges arise ^8,9,10^.

It has been suggested that the efficiency of some enzymes is affected by the kinetics of the substrate binding or product unbinding. This may happen if small molecule diffusion along these routes become rate limiting e.g. when the small molecule, in its interaction with the protein, assumes a particular energetically unfavorable configuration. These regions in protein tunnels are called bottlenecks, and they are clearly hotspots for mutagenesis ^11,12^. Direct observation of a ligand binding process and bottleneck identification is impossible with current experimental methods, and researchers resort to computational simulations. One approach here is to use molecular dynamics simulations with enhanced sampling, to study possible one-way unbinding events (as in τ-Random Acceleration Molecular Dynamics ^13,14^) or sample instances of a complete reversible process (as in funnel metadynamics ^15–17^). These techniques are computationally demanding and thus are performed for singular cases. As a result, not only experimental, but also synthetic datasets and benchmark sets of “ground truth” protein-ligand binding processes do not exist. Despite being highly physically adequate, these MD-based approaches are also too computationally expensive to serve within design pipelines as filters to assess and rank the properties of (re)engineered enzymes. Nevertheless, a demand for such capabilities has been actively discussed in the community ^18^.

Simpler computational approaches are being developed to study the ligand (un)binding process based on small molecule docking ^19^. These approaches are based upon a crucial conceptual assumption that might not necessarily be true for a corresponding real-life process, but makes the task formalization tangible. The process, which is time-resolved in reality, is modeled as a discrete and strictly monotonic progression in space. Possible backtracking events are therefore not accounted for, which is equally true for steered MD simulations, sometimes used in the field ^20^. Nevertheless, these approaches gained recognition and are used in practice due to their low computational cost and uncomplicated set up. Currently available tools include MoMA-LigPath ^21^, GPathFinder ^22^, and a highly popular CaverDock ^23^. They estimate an optimal ligand pose for each point along the process independently, which might pose further challenges. Significant changes in a ligand conformation happening under small displacements could be considered unphysical, hampering the interpretability of the results. Therefore, being able to control conformational continuity emerges as a practically important feature. CaverDock addresses this challenge at a post processing step, and by rerunning calculations. This, however, makes runtimes unpredictable, and for some systems a continuous solution may still not be found ^23^. Moreover, while CaverDock models the ligand fully flexible, the protein is treated statically, with side chain rotamers fixed. This limitation can be alleviated by repeating the calculations with multiple different protein input conformations – however, this would again extend the runtime. We therefore hypothesized that a computational approach that is explicitly designed to provide conformational continuity and protein flexibility within a single run could be potentially useful for the community and complement existing tools.

Rosetta is a well-established software suite for protein modeling and design ^24^. Extensive development put RosettaLigand within the premiere solutions for small molecule docking ^25^, owing to its full protein and ligand flexibility and scoring function. Access to its capabilities through a widely customizable and extensively documented RosettaScripts ^26^ XML interface gives the user full control of the ligand docking protocol ^27^. In this work, these features empowered us to build a computational approach to study protein-ligand (un)binding pathways that directly meets the aforementioned requirements of continuity and flexibility. The resulting engine – RosettaLigand PathFinder (RLPF) – ensures that every run finishes with an interpretable output under predictable compute requirements comparable to those of other docking-based tools. RLPF is made highly customizable to cater for various use scenarios. We demonstrate the utility of the tool in a number of cases, showing its power to differentiate between ligands and pathways, identify bottlenecks and known secondary binding sites, and correctly capturing the known effects of mutations.

## Results and Discussion

### RosettaLigand PathFinder is designed to capture physically realistic (un)binding trajectories

We present here RosettaLigand PathFinder (RLPF), a tool that builds upon Rosetta to sample ligand (un)binding trajectories (Figure 1A). Its inputs are a) a pdb file of a protein-ligand complex in the starting conformation, b) a guide path, and c) input parameters (see Methods and SI). A guide path is a set of points in 3D space that, after spline interpolation, is used to specify a particular direction of (un)binding. This way, for example, an exploration of a particular tunnel or direction can be requested, if multiple tunnels are possible. To generate the guide path, fast ligand-agnostic tools that rely on the geometry of the protein can be used, including CAVER ^28^, MOLE ^29^, and AQUA-DUCT ^30^. Alternatively, we provide a slower but ligand-aware routine built with Rosetta (see SI). Finally, the path can be set up with just a handful of points chosen manually, which can be heRLPFul in a general case with no tunnels. This way, multiple input options are provided to align with various use scenarios. During its (un)binding path exploration, RLPF allows for different setups concerning the influence of the guide path on the modeling. A precise following of the path might be requested, alongside more relaxed scenarios with only a general alignment in direction, all the way to almost free roaming (Figure 1E). Finally, a list of considerations includes protein flexibility, allowing for sidechain movements, rotamer changes, and small backbone adjustments. The degree of protein flexibility is also freely adjustable through XML protocols.

**Figure 1.**
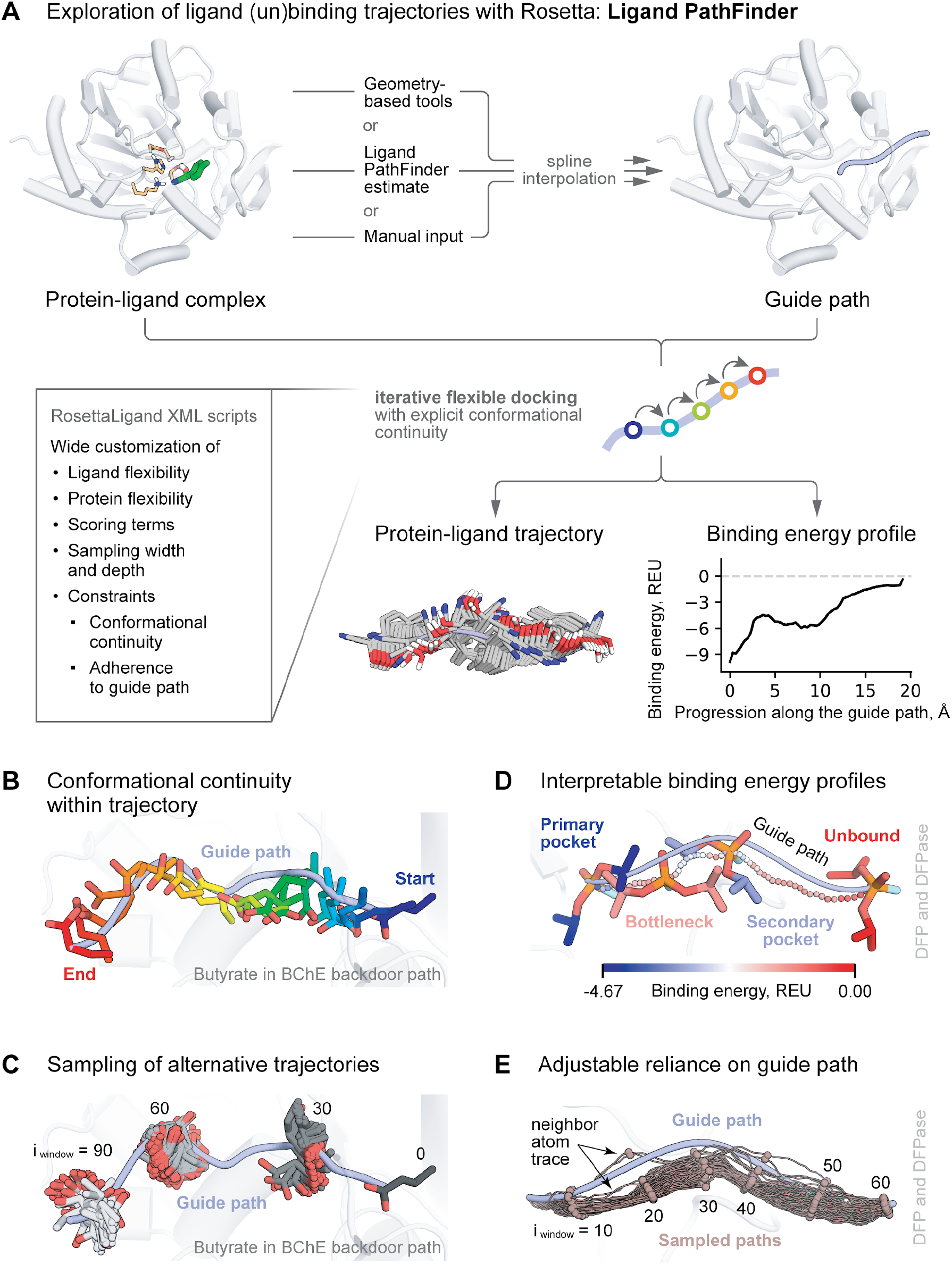
An overview of RosettaLigand PathFinder (RLPF) algorithm and features. **A**. A workflow of RLPF, including compulsory inputs, outputs, and main adjustable parameters. **B**. An example of an RLPF trajectory with explicit control over ligand conformational continuity. Only selected snapshots along the path are shown for clarity. **C**. A demonstration of the diversity of independently sampled paths. Shown are ligand conformations assumed at 30th, 60th, and 90th iterations of RLPF. **D**. An example of ligand neighbor atom trace colored by binding energy show biophysical interpretability of modeled states. **E**. A demonstration of relative adherence of trajectory sampling to the guide path, the degree of which can be precisely controlled by the user.

As its core logic, RLPF performs a series of conformationally and positionally constrained docking simulations. For that, well-established RosettaLigand protocols for flexible docking are used, expressed through a highly customizable XML RosettaScripts interface. This way, we provide a wide range of adjustable parameters to the user (see Methods and SI). For each run, RLPF outputs protein-ligand trajectories in PDB format and binding energy profiles in Rosetta Energy Units (REU).

RLPF was built to satisfy a number of practical considerations that differentiate it from similar tools. First and most important is an explicit control over conformational continuity (Figure 1B). Through adjustable restraints (termed “constraints” in Rosetta internally), it is ensured that each run outputs a trajectory without unphysically large differences in ligand pose and conformation upon small incremental advancements along the (un)binding path. RLPF logic ensures that a solution is found in every run, as compared to a more uncertain behavior of CaverDock (see SI).

Second, RLPF is envisioned to be used in an ensemble regime, benefiting from the inherent stochastic nature of Rosetta sampling. Comparably low computational requirements (see Methods) allow for sampling dozens or even hundreds of trajectories (Figure 1C) to then analyze them statistically or perform clustering. This results in statistically robust and reproducible inference of biologically relevant insights (Figures 2-5).

**Figure 2.**
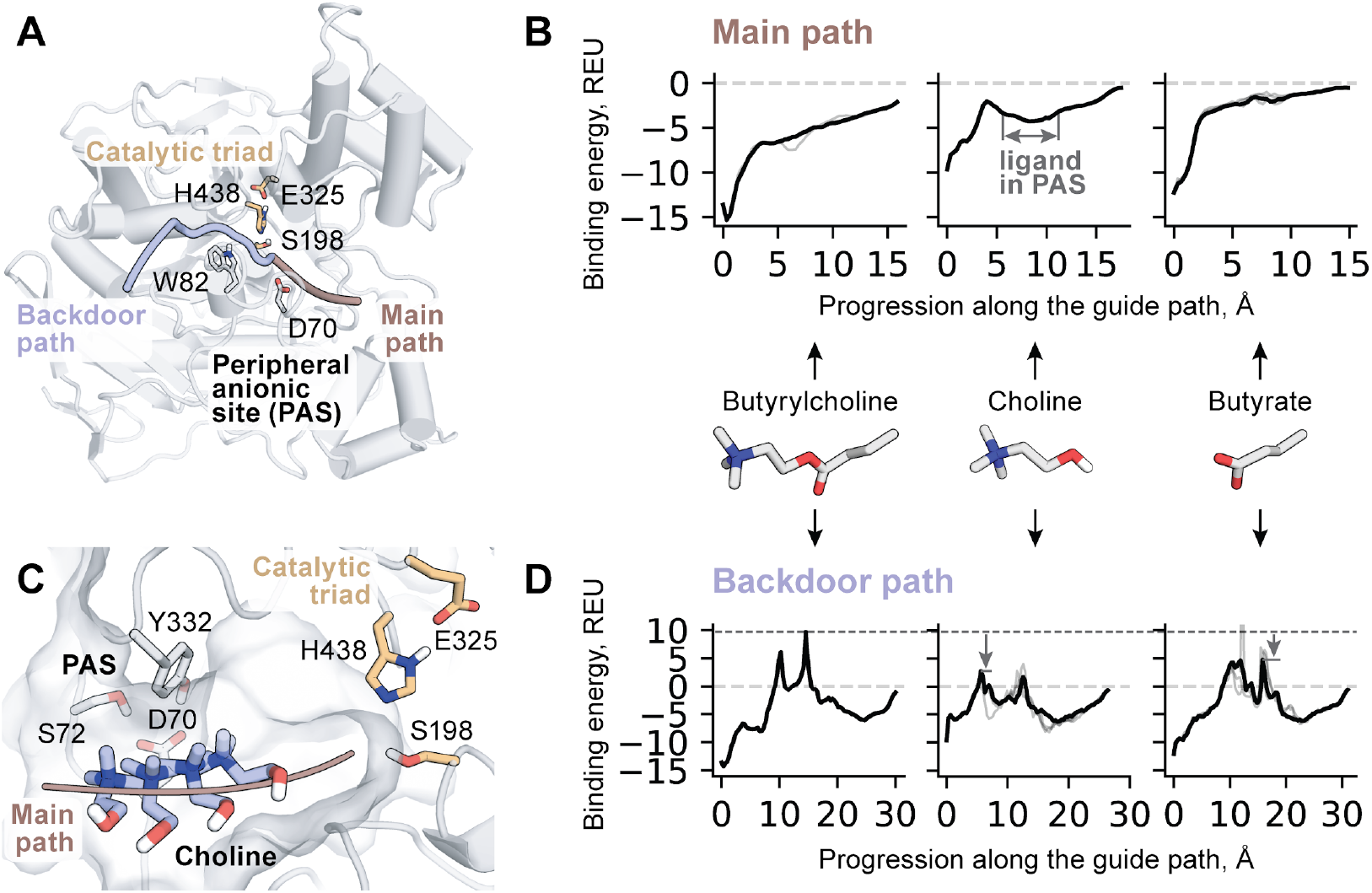
Analysis of ligand unbinding through main and backdoor tunnels of human butyrylcholinesterase. **A**. Overview of the enzyme structure and guide paths used in the study. **B**. Binding energy profiles for the main path show an obscured unbinding for butyrylcholine and butyrate, and a secondary stable binding mode of choline in the peripheral anionic site (PAS). **C**. Selected snapshots along the unbinding trajectory of choline when it is interacting with PAS. Only one snapshot is used to show enzyme atoms for clarity. **D**. Binding energy profiles for the backdoor path show differences in the probability of passing the bottleneck. **B, D**. A thick black line represents the mean energy of the most populated trajectory cluster, gray lines correspond to the means of minor clusters.

RLPF outputs a binding energy profile for each trajectory, connecting a particular state along the (un)binding pathway to its binding energy (Figure 1D). This energy is calculated with Rosetta scoring functions ^31^, with precise control over any terms allowing for highly customized setups. While REU relation to any physical units is not direct, relative differences in Rosetta-calculated energies are interpretable ^32^ and point to physically relevant insights ^33^. This way, analysis of RLPF-generated profiles can help mapping various stable bound poses and bottlenecks along (un)binding paths and understand the rationale for them.

The intended use of RLPF is to sample and analyze multiple conformationally constrained trajectories for each input with explicit protein and ligand flexibility. These features differentiate it from similar tools and prevent direct comparison. While it is technically possible to imitate their behavior by disabling protein flexibility and conformational continuity, it is not an intended setup. Therefore, we believe RLPF to be a valuable and orthogonal addition, rather than competition, to the family of ligand (un)binding tools.

### RosettaLigand PathFinder produces biologically relevant insights

To showcase the capabilities and scope of RLPF, we applied it to a number of enzymatic systems with known properties tied to the ligand (un)binding.

### Differentiating between pathways and ligands

For human butyrylcholinesterase, an existence of a transient backdoor has been previously suggested ^34^. Through this backdoor (Figure 2A), smaller products of the reaction are thought to be able to pass, and not a larger substrate ^35^. All compounds were shown to pass well through the main tunnel. A prominent feature of the main tunnel is a peripheral anionic site (PAS) centered at Asp70 ^36^. This site attracts cationic ligands and is thought to be crucial for substrate activation ^37^.

We performed RLPF calculations for both pathways and three ligands – a substrate, butyrylcholine, and both products of the reaction, choline and butyrate. We used CaverWeb to generate the initial guiding paths (see Methods). For the main path, RLPF yielded smooth and funnel-shaped binding energy profiles for butyrylcholine and butyrate (Figure 2B) that never reached into the positive (unfavorable) binding energy ranges. This means that the passage along this path is unhindered for these ligands, and tunnel-lining residues promote the diffusion towards the catalytically competent binding pose. For a positively charged choline, the PAS was uncovered (Figure 2C) as a -4.3 REU local minima, in agreement with what is known about the enzyme. A small bottleneck was found between the PAS- and the active site-bound poses of choline as indicated by an energy peak roughly 4.5 Å away from the active site. This state, however, still had a below-zero binding energy of -2.1 REU. In contrast, analysis of ligand transport through the backdoor revealed above-zero energy bottlenecks for all three ligands. This way, RLPF clearly established the main pathway as more preferred. Analysis of ligand transport through the backdoor revealed the lowest bottleneck of +2.7 REU for a product choline, followed by butyrate with +4.8 REU and then a substrate, butyrylcholine (Figure 2D) with +9.7 REU, in agreement with the suggestion of a previous study ^35^.

While RLPF treats proteins as flexible, this flexibility mostly concerns side chains. Above-zero binding energies along the backdoor path, calculated with a particular protein conformation used in the study, might point to the induced-fit nature of this tunnel, with significant backbone shifts happening to allow the passage in reality. Sampling such conformational changes is currently beyond the scope of RLPF, but can nevertheless be achieved by using multiple protein conformations at input. Methods of modeling such conformational ensembles ^38–40^ are beyond the scope of this work. We note that the same consideration regarding positive binding energy values apply to all other systems studied in this work.

The second enzyme we studied was diisopropyl fluorophosphatase (DFPase) with its substrate DFP. We have previously performed microsecond-scale funnel metadynamics simulations to map the enzyme-substrate binding-unbinding process in great detail ^16^. We showed that the pre-reactive pose is reached by first binding at a secondary binding site on the right side of the pocket. Here, we used RLPF to reproduce these findings in a fraction of time that had been previously required (Figure 3). We modeled two unbinding directions based on CaverWeb-generated paths, leaning to the left and to the right sides of the binding pocket respectively. RLPF binding energy profiles for the left path featured a +3.4 REU bottleneck 5.9 Å away from the pre-reaction pose. A much more modest, negative energy bottleneck of -1.2 REU at 5.3 Å was found for the right path, establishing it as a preferred access route (Figure 3B) in agreement with our previous results. A local minimum -2.4 REU deep, 6.9 – 12.2 Å away from the active site corresponded well to a previously identified secondary binding site (Figure 3C).

**Figure 3.**
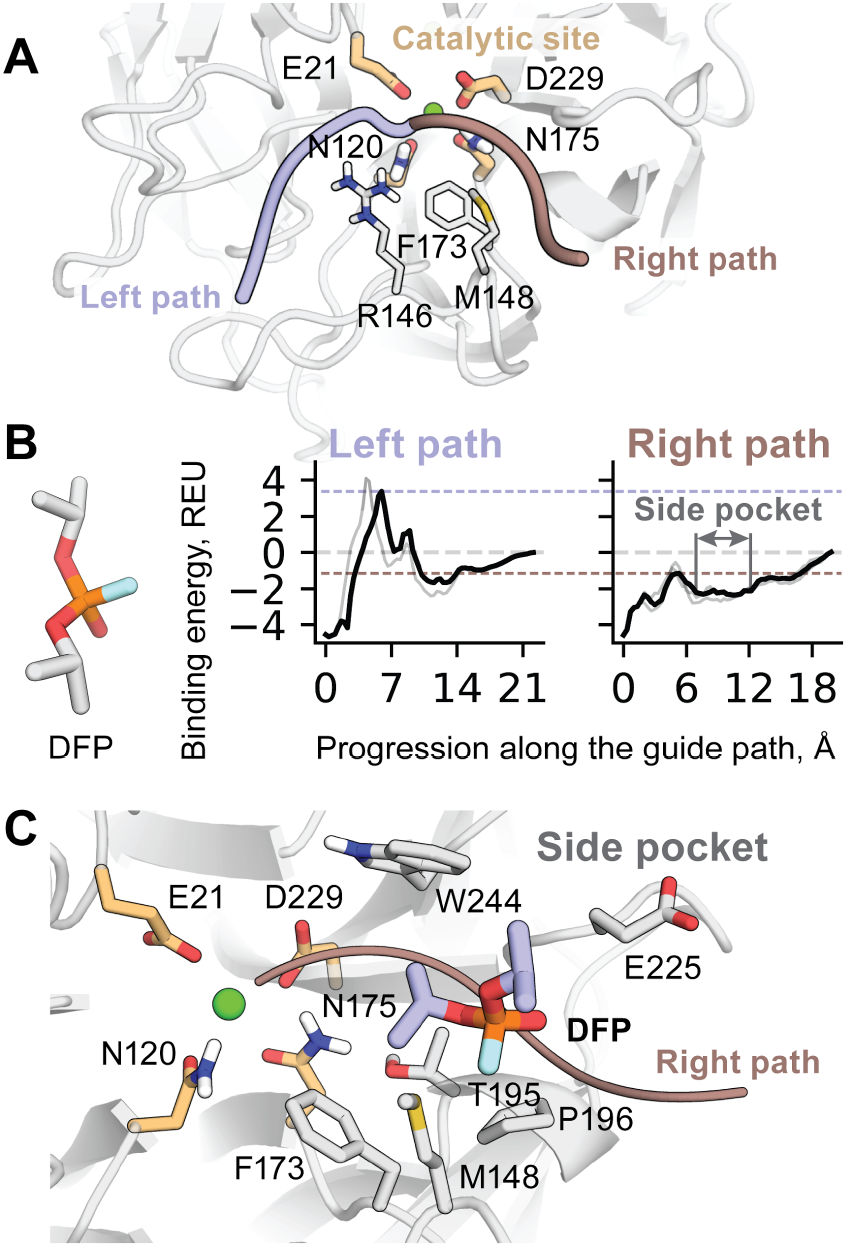
Analysis of ligand unbinding of diisopropyl fluorophosphatase substrate. **A**. Overview of the enzyme structure and guide paths used in the study. **B**. Binding energy profiles when substrate is guided through the left and right side of the pocket. A thick black line represents the mean energy of the most populated trajectory cluster, gray lines correspond to the means of minor clusters. **C**. An example of a side pocket-bound pose modeled by RosettaLigand PathFinder, corresponding closely to previously reported results (^16^).

Next, we studied two haloalkane dehalogenases with dramatically different substrate preferences and tunnel shapes – DhlA and DmmA. It was shown that the former prefers smaller substrates like ethylene dichloride (EDC), while the latter prefers larger ones including 4-bromobutyronitrile (4Br-BN) ^41,42^. We applied RLPF to them and reproduced these trends (Figure 4). Binding energy profiles for DhlA were found to feature a substantially lower bottleneck for EDC passage than for 4Br-BN (Figure 4C), with peaks of +5.8 and +18.4 REU respectively. Profiles calculated for DmmA, in contrast, were absent of bottlenecks, were smooth and funnel-shaped (Figure 4D). Bound poses of a larger ligand had lower REU scores (thus, were bound better) than those of a smaller ligand, with -4.0 and -2.4 REU respectively. Thus, the analysis of energy profiles allowed us to hypothesize that the preference for small ligands in DhlA originates from kinetic effects because of the lower modeled energy barrier at the bottleneck. At the same time, preference for larger substrates in DmmA might have a thermodynamic nature (Figure 4D), with no detectable bottleneck due to a larger tunnel radius ^41,42^.

**Figure 4.**
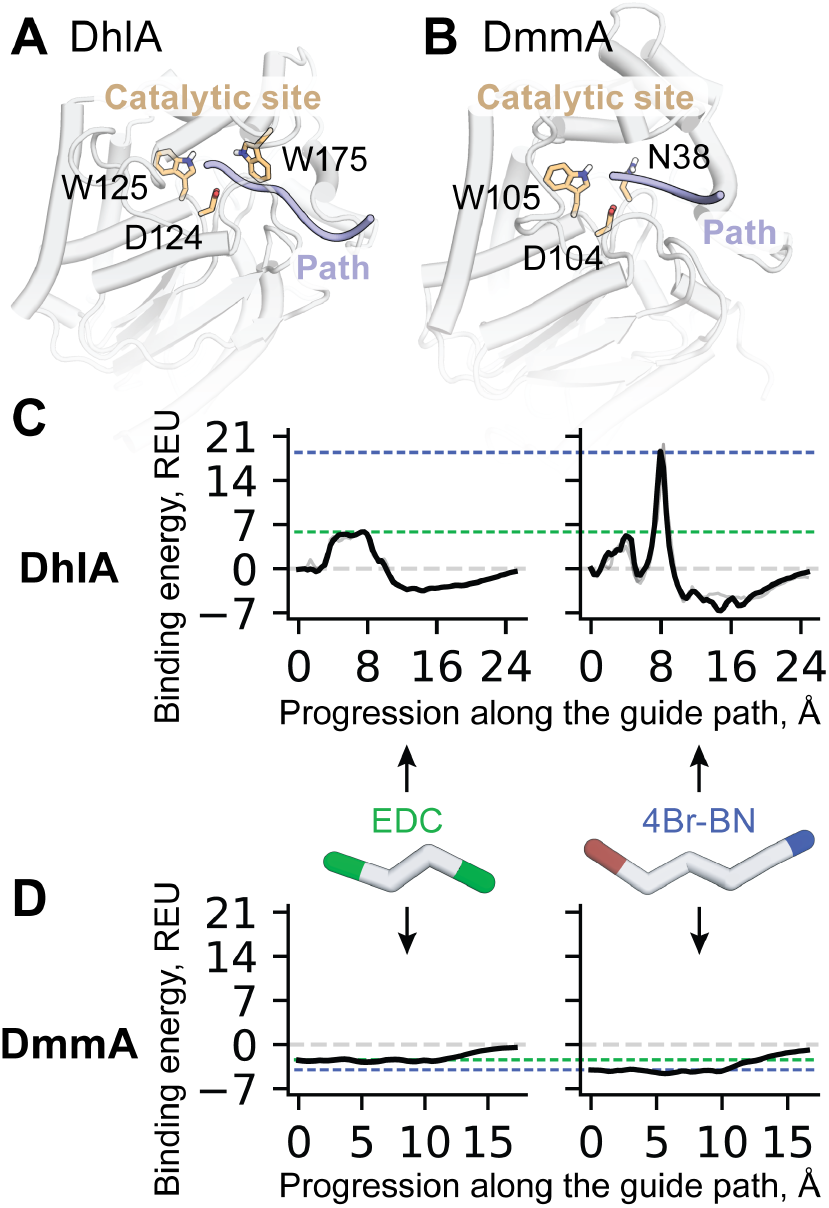
Analysis of ligand unbinding from haloalkane dehalogenases DhlA and DmmA. **A**. An overview of DhlA, its catalytic site, and a guiding path used in the study. **B**. An overview of DmmA, its catalytic site, and a guiding path used in the study. **C**. Binding energy profiles for EDC and 4Br-Bu substrates passing through the tunnel in DhlA. **D**. Binding energy profiles for substrates passing through the tunnel in DmmA. A thick black line represents the mean energy of the most populated trajectory cluster, gray lines correspond to the means of minor clusters.

This way, we demonstrated that RLPF can reproduce established transport characteristics across several well-studied enzymes within a consistent computational framework. The method proved sensitive enough to distinguish alternative pathways and ligand-dependent preferences. We suggest that, beyond reproducing known behavior, RLPF can be used to rapidly generate hypotheses on fundamental properties and structure–function relationships of enzymes, providing seed ideas for subsequent large-scale research campaigns. As an example, it can be used to analyze the presence of eventual diffusion bottlenecks, identify the nature of such bottlenecks, identify the amino acids responsible and nominate hotspots for mutagenesis.

### Capturing effects of mutations in tunnel-forming residues

We then investigated how RLPF performs for cases, when experimentally measured changes in kinetics were observed upon a single amino acid substitution. We carefully selected three systems for which the chances that these structure-function associations were tied to the ligand binding process – as opposed to the efficiency at the chemical step, or binding affinity – were deemed substantial.

For a haloalkane dehalogenase DhaA31 (Figure 5A), it has been shown that Y176A substitution improves activity on a substrate 1-chlorohexane ^43^. It has been suggested that the reason is a better passage through the bottleneck. We applied RLPF to build binding energy profiles along the (un)binding path of this ligand from both the WT and Y176A variants of the enzyme (Figure 5D). Most of the trajectories for the WT were modeled with binding energies well above zero REU. In comparison, with Y176A we observed a dramatic -39.1 REU reduction in the energy level of the highest bottleneck along the path, bringing its absolute value to +0.5 REU, close to zero. This indicates a much less obscured passage in this conformation and might point to a higher energetic cost for a WT to adapt to a more open tunnel conformation via induced fit in reality.

**Figure 5.**
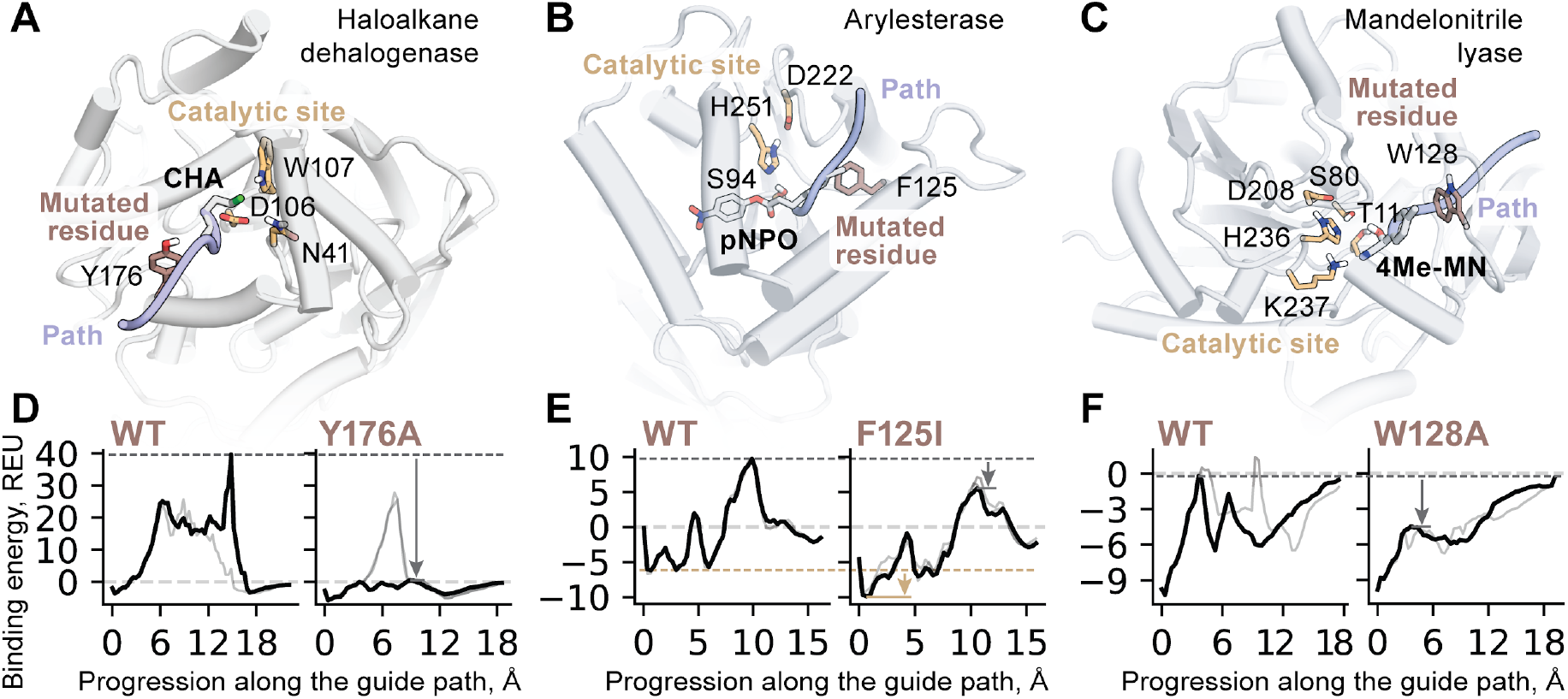
Analysis of ligand unbinding uncovers effects of mutations in tunnel-forming residues. **A**. An overview of haloalkane dehalogenase DhaA31, its catalytic site and substrate, a guiding path used in the study and a substituted residue of interest. CHA = 1-chlorohexane, a substrate. **B**. An overview of arylesterase. pNPO = 4-nitrophenyl octanoate, a substrate. **C**. An overview of mandelonitrile lyase. 4Me-MN = 4-methyl mandelonitrile, a substrate. **D**. Binding energy profiles for the substrate passing through the tunnel in DhaA31. **E**. Binding energy profiles for the substrate passing through the tunnel in arylesterase. **F**. Binding energy profiles for the substrate passing through the tunnel in mandelonitrile lyase. **D-F**. A thick black line represents the mean energy of the most populated trajectory cluster, gray lines correspond to the means of minor clusters.

For arylesterase from Pseudomonas fluorescens (Figure 5B), substitution of a tunnel-lining bulky F125 for a smaller Ile was shown to result in an order of magnitude higher activity ^44^. We again subjected this system to the analysis with RLPF. We observed that the bottleneck energy value dropped by -4.2 REU with the introduction of the F125I substitution, while the pre-reactive pose was also stabilized by -3.8 REU. These findings suggest that F125I potentially brought both an improved affinity and a relaxed bottleneck (Figure 5E) as reasons for the improved activity.

Similarly, mandelonitrile lyase activity was found to improve upon substitutions of a bulky W128 at the entrance to the binding pocket (Figure 5C) ^45^. We modeled the unbinding trajectories of a substrate, 4-methyl mandelonitrile (4Me-MN), with RLPF. While profiles for both WT and W128A featured a local maximum 3.6 Å away from the active site, its energy in W128A was -4.5 REU, dropping from -0.2 REU in WT (Figure 5F). Such a difference means that the bottleneck for substrate binding is sufficiently relaxed in W128A, aligning with the improvement in activity.

The results presented in this section show that RLPF could be potentially useful as a robust tool for ranking of enzyme variants in engineering campaigns based on their tunnel and ligand binding kinetics properties.

## Methods

### RosettaLigand PathFinder algorithm

RLPF is a Python application that serves as a wrapper to RosettaScripts ^26^. The wrapper performs calculations and file management, preparing sequential RosettaLigand ^27^ docking runs. Its inputs are arranged in a single json file that contains both paths to necessary files and run parameters. Files necessary to initiate a run include the starting pdb file of the protein-ligand complex, the guide path, and a folder with typical Rosetta-generated ligand parameters (see protocol capture in SI and example files).

At the start of the calculation, an input guide path is smoothed by spline interpolation, up-sampled and then discretized to a set of guide points with a user-defined step size (in Å) between consecutive points. These points and the local curvature are used to calculate the associated normal vectors to define planes passing through each point orthogonal to the path direction. A RosettaLigand docking run associated with a single particular plane is referred to as a “window”. In such a run, the location of the ligand is kept tied to the respective plane. These runs are performed sequentially for all the points of the discretized path, with the following run depending on the results of the previous one, in contrast to an independent approach used in CaverDock ^23^. Docking is performed according to a user-specified XML protocol (with templates provided in the supplementing files) in a number of replicates, and the best pose is saved as a solution of the window. The selection of the best pose can be set based on the total score of the whole system or on the protein-ligand interaction score. The final trajectory, therefore, relates to a set of such solutions from all windows.

A set of restraints is constructed by the wrapper and passed to the docking cycle to achieve that ligand stays associated with the plane, and its conformation and location are similar to the solution in the previous windows (Figure 6, Figure 1). The latter is central to the RLPF design objective of explicitly providing conformational continuity of the trajectory.

**Figure 6.**
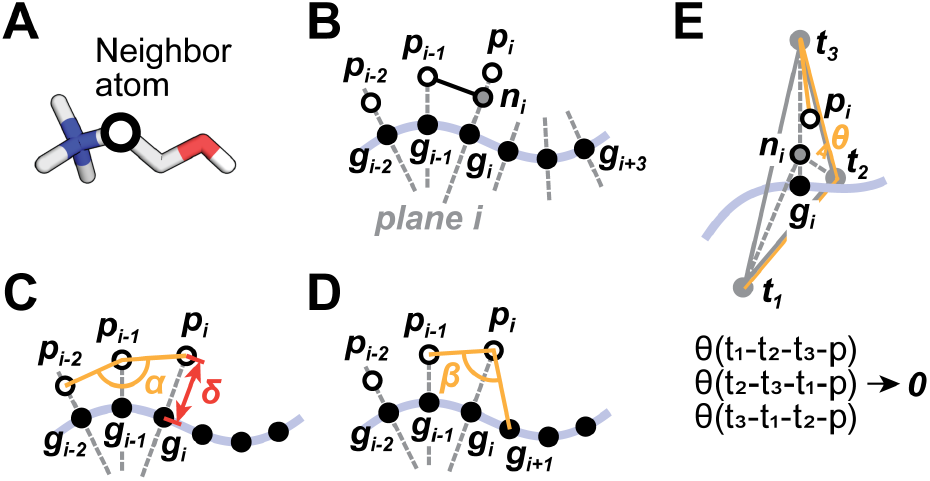
Notations and restraint types used in RosettaLigand PathFinder. **A**. Restraints are applied to the location of the neighbor atom, typically the most central atom in the molecular graph. **B**. Each iteration starts from the projection (n_i_) of the previous result (p_i-1_) onto the next plane defined by the guide path, and solutions are searched for within this plane. **C**. The angle α between the neighbor atom and its positions in two previous solutions can be controlled. Additionally, a maximum orthogonal displacement from the guide path (δ) can be specified. **D**. An angle β between the previous solution, a current location of the neighbor atom, and the next point on the guide path can be controlled to prevent cumulatively divergent exploration. **E**. An in-plane restraint is implemented by three torsion restraints built with the vertices of an equilateral triangle centered at point n_i_.

One ligand atom is used to control its translational and rotational motion as a whole. As is typical in Rosetta, here we use a neighbor atom, which is automatically assigned by Rosetta parameterization utilities as a central-most atom in the molecular graph, but can also be redefined by the user (Figure 6A). A calculation in the window starts by finding a projection of this atom to the new plane, and shifting the whole ligand molecule on the resulting vector (Figure 6B). This way, the neighbor atom now lies precisely in the current plane. Constraining the ligand to the current plane is achieved through a combination of three torsion restraints (Figure 6E) applied to the location of the neighbor atom. Restraints that ensure the continuity of a ligand trajectory include angular (Figure 6C) and in-plane displacement restraints (Figure 6D), which are also resolved at a neighbor atom level, and an RMSD restraint, acting on all (or, optionally, all heavy) atoms of the ligand. For RMSD restraint, reference coordinates of the ligand atoms are the starting coordinates of this window before the docking run, but after the ligand has been shifted so its neighbor atom lies within the plane. This is done to make the influence of this setting independent of the selection of the step size (as opposed to using the solution in a previous window as reference). All listed restraint potentials can be specified as either harmonic or flat-bottomed with arbitrary strengths to achieve full control over the process. We make use of restraint potentials already available in RosettaScripts. All restraints except for keeping the ligand tied to a plane are optional.

A full list of adjustable run parameters can be found in the SI, alongside the tutorial for RLPF. We provide recommended starting values for all restraints and other run parameters in the run files for the systems used to illustrate RLPF capabilities in the paper.

### Modeling details

AlphaFold3 ^46^ was used to provide initial coordinates for enzyme-substrate complexes. These poses were manually assured to correspond to possible pre- and post-reaction states. Then ligand coordinates were transplanted into experimental models from PDB (see SI) and refined with flexible RosettaLigand docking. Using the BioChemical Library (version 4.3.0) ^47^, we generated up to 1000 conformers for each compound, clustered by RMSD with a threshold of 0.25 Å. Guide paths were obtained with CaverWeb ^42^ and extended 5-10 angstroms to better sample unbound poses. For each enzyme-ligand-path system, we sampled 100 independent trajectories with RLPF, and clustered them with the AffinityPropagation algorithm (see SI for details). Throughout the paper, only clusters with more than 10 samples are retained for analysis, and average binding energies are calculated for each window within a cluster. We rely on averaging to counteract the intrinsically stochastic nature of Rosetta sampling and approximate nature of solutions within each window. Rosetta docking and RLPF XML protocols, parameters and input files, including guide paths, are provided with the manuscript.

## Conclusions

In this work, we introduced RosettaLigand PathFinder (RLPF), a protocol that leverages the Rosetta framework to model ligand (un)binding pathways under explicit control of conformational continuity and protein flexibility. RLPF bridges the gap between rigid geometric tunnel analysis and computationally intensive molecular dynamics approaches, providing a consistent and reproducible means to explore ligand motion in proteins, including enzymes. RLPF principles and parameters would be familiar to any experienced Rosetta user. Applied to a range of well-characterized systems, RLPF accurately captured known transport routes, bottlenecks, and effects of mutations on tunnel properties, highlighting its ability to produce biologically relevant insights.

A key strength of RLPF lies in its customizability. Built upon the well-established and extensively documented RosettaScripts XML interface, it enables fine-grained control over scoring terms, sampling parameters, restraint potentials, and degrees of protein and ligand flexibility. This design allows users to adapt the protocol to diverse use cases. RLPF is written in Python and is freely available. In developing RLPF, we deliberately prioritized conformational continuity, ensemble sampling, and explicit protein flexibility. These choices define its practical scope and make it a complementary addition to existing tools for studying ligand transport. We envision that RLPF can be used both in fundamental and applied settings. In exploratory studies of proteins it might help formulate structure-function hypotheses, while in enzyme engineering campaigns it can prioritize a starting enzyme or its mutant variant from a list of candidates based on the transport properties of a substrate, a product or an inhibitor.

## Data and Code Availability

Rosetta docking and RLPF XML protocols for all enzyme-ligand systems, parameters and input files, including guide paths, are provided in a Zenodo repository 10.5281/zenodo.23183177.

The code will be made available at GitHub at https://github.com/meilerlab/LigandPathFinder.

## Acknowledgements

J.M. was supported by an Alexander-von-Humboldt-professorship from the Alexander-von-Humboldt foundation and by the Federal Ministry of Education and Research of Germany and by the Sächsisches Staatsministerium für Wissenschaft Kultur und Tourismus in the program Center of Excellence for AI-research “Center for Scalable Data Analytics and Artificial Intelligence Dresden/Leipzig”, project identification number ScaDS.AI.

A.Z. and J.M. acknowledge funding by the Federal Ministry of Education and Research of Germany (BMBF) in the frame of de.NBI/ELIXIR-DE (Grant/FKZ: W-de.NBI-018).

