## Supplementary Information for "Exploration of ligand (un)binding trajectories with Rosetta"

### Contents

**Built-in guess path estimation (coarse_path algorithm)** 3
**Geometry of Tunnels benchmark** **4**
Description 4
Reference 4
Run times 4
**Input JSON configuration reference** **10**
Top level 10
input 10
parameters 11
Path geometry 11
Conformer sampling 11
RMSD (shape) constraint 11
In-plane-deviation constraint (ipdc) 12
Plane (coplanarity) constraint 12
Direction (angle) constraints 13
Pose propagation & selection 13
Optional pre-docking 13
rosetta_scripts 14
parameters.run_stats / statistics pass 14
output 15
Outputs written to output.path 15
**Protocol capture** 16
Requirements 16
1. System preparation 16
1.1 Binding pose 17
1.2 Rosetta parameters for the ligand 17
1.3 Starting file for Ligand PathFinder 19
2. Coarse path finding 20
2.1 Preparation 20
2.2 Run 22
3. Trajectory 24
4. (Optional) Replicates 27
5. Adjustments 28

### Built-in guess path estimation (coarse_path algorithm)

When no guide path is available, RLPF can generate one with the coarse-path routine. The input consists of two points: the position of the neighbor atom of the ligand in the bound pose and an endpoint in the solvent, which roughly defines the direction of unbinding. Optionally, the endpoint is displaced within a disc orthogonal to the line connecting the two points, so that repeated runs sample different endpoints and produce an ensemble of paths that can be clustered.

The path is constructed by recursive bisection. The first point is placed at the middle of the segment between the two input points. For each point, the ligand is shifted from the structure of the adjacent solved point so that its neighbor atom coincides with the target position, and a RosettaLigand docking run is performed in a user-specified number of replicates. The best pose, selected by the total score of the system or by the protein-ligand interaction score, is saved as the solution of this point, and the position of its neighbor atom is taken as the coordinate of the path point. The in-plane restraint, similar to one from the main algorithm, is active at all times. Two angular restraints keep the ligand between its neighbors on the path and prevent the path from folding back, and an RMSD restraint keeps the ligand conformation close to the starting structure of the run, with a width that increases with the distance to the starting position. All restraints are specified as harmonic or flat-bottomed potentials with user-defined strengths, and, except for the plane restraint, are optional.

After a point is solved, the distances to its left and right neighbors are evaluated. If a distance exceeds the user-defined step size, a new point is inserted at the middle of the corresponding gap and solved in the same way. The procedure is repeated until all distances between consecutive points do not exceed the step size, and the number of points therefore depends on the length of the path. The ordered neighbor atom coordinates, starting from the bound pose, form the coarse path, which is used as the guide path for the trajectory calculation.

### Geometry of Tunnels benchmark

#### Description

To test the robustness of the RLPF trajectory protocol, we applied it to the d2_Geometry_of_tunnels dataset introduced with CaverDock (Vavra et al., 2019; see Reference). The dataset comprises 136 cases (protein, tunnel and native substrate combinations) covering a broad range of tunnel geometries and ligand sizes. As it stands, the dataset does not guarantee that the tunnels are easily traversable by the ligands, with or without large-scale protein-side readjustments, which makes it a good test of robustness: the aim was to verify that the protocol reaches the end of the path regardless of the technical details of a run and of the possible unphysicality of a given ligand, protein and tunnel combination. We also used this benchmark to estimate the run time of the protocol. The ligands had on average 41 atoms (19 heavy atoms; range 4-159 and 2-59, respectively), and the average run time was 2.7 min per frame (SD 1.0 min; range 0.9-8.2 min) on a single CPU core. All 136 systems ran to technical completion. For comparison, CaverDock completed 125 of the 136 cases (92%) with its lower-bound trajectory, and 81 (60%) with its upper-bound trajectory; in 11 cases the ligand was unable to pass through the tunnel. Our approach has no separate lower- and upper-bound modes, and its results are more directly comparable with the upper-bound trajectories.

Shared run settings:

- The original CAVER path was used as a guide path.
- Path step 0.5Å and smoothing 5.0 were used.
- 10 structures per frame, with the best one chosen by total_score. That structure is also the start for the next frame.
- Constraints:
    - harmonic RMSD constraint with sd 0.66Å
    - in-plane deviation constraint ($d_0$ 2.0Å, sd 0.5Å, plane sd 0.02 rad.)
    - angle constraints (bound 2.5 rad., sd 0.5 rad., and bound 1.57 rad., sd 0.25 rad. for the next-frame angle)
- Conformer subsampling threshold 3.0Å and outlier RMSD 2.0Å for normal-sized ligands, 5.0Å and 5.0Å respectively for excessively large ones (more than 50 atoms).
- One serial run on one CPU core per system.

Input and configuration files for all benchmark runs are shared on zenodo:

### Run times

Total time is the wall time to build the full trajectory for a tunnel. Path length is the number of frames. All runs were performed in a single-core regime with no parallelization.

| part | protein | tunnel | path length | ligand atoms | ligand heavy atoms | total time | avg time/frame |
|---|---|---|---|---|---|---|---|
| part1 | 1AKD | CD01 | 26 | 27 | 11 | 58m 45s | 2m 15.58s |
| part1 | 1AKD | CD02 | 29 | 27 | 11 | 1h 15m 23s | 2m 35.97s |
| part1 | 1AKD | CD03 | 27 | 27 | 11 | 1h 3m 26s | 2m 20.96s |
| part1 | 1MQF-2 | CD01 | 28 | 4 | 3 | 50m 15s | 1m 47.68s |
| part1 | 1MQF-2 | CD02 | 21 | 4 | 3 | 34m 29s | 1m 38.52s |
| part1 | 1MQF-2 | CD03 | 39 | 4 | 3 | 59m 32s | 1m 31.59s |
| part1 | 1MQF | CD01 | 28 | 4 | 2 | 41m 56s | 1m 29.86s |
| part1 | 1MQF | CD02 | 21 | 4 | 2 | 32m 2s | 1m 31.52s |
| part1 | 1MQF | CD03 | 39 | 4 | 2 | 57m 16s | 1m 28.10s |
| part1 | 1MXT | CD01 | 31 | 74 | 28 | 1h 58m 47s | 3m 49.90s |
| part1 | 1MXT | CD02 | 33 | 74 | 28 | 2h 38m 3s | 4m 47.36s |
| part1 | 1MXT | CD03 | 57 | 74 | 28 | 3h 43m 38s | 3m 55.40s |
| part1 | 1THG | CD01 | 18 | 159 | 59 | 1h 10m 8s | 3m 53.78s |
| part1 | 1THG | CD02 | 20 | 159 | 59 | 1h 25m 52s | 4m 17.60s |
| part1 | 1THG | CD03 | 25 | 159 | 59 | 1h 20m 39s | 3m 13.56s |
| part1 | 2ACE | CD01 | 22 | 26 | 10 | 1h 3m 13s | 2m 52.41s |
| part1 | 2ACE | CD02 | 19 | 26 | 10 | 52m 38s | 2m 46.21s |
| part1 | 2ACE | CD03 | 30 | 26 | 10 | 1h 38m 0s | 3m 16.00s |
| part1 | 2BG9 | CD01 | 265 | 26 | 10 | 19h 39m 45s | 4m 27.11s |
| part1 | 2JBV | CD01 | 37 | 21 | 7 | 1h 35m 41s | 2m 35.16s |
| part1 | 2JBV | CD02 | 52 | 21 | 7 | 2h 3m 55s | 2m 22.98s |
| part1 | 2JBV | CD03 | 43 | 21 | 7 | 1h 42m 33s | 2m 23.09s |
| part2 | 1A9X-2 | CD01 | 19 | 5 | 4 | 32m 49s | 1m 43.63s |
| part2 | 1A9X-2 | CD02 | 44 | 5 | 4 | 1h 14m 18s | 1m 41.32s |
| part2 | 1A9X-2 | CD03 | 40 | 5 | 4 | 55m 47s | 1m 23.67s |

| part | protein | tunnel | path length | ligand atoms | ligand heavy atoms | total time | avg time/frame |
|---|---|---|---|---|---|---|---|
| part2 | 1A9X-3 | CD01 | 19 | 20 | 10 | 55m 5s | 2m 53.95s |
| part2 | 1A9X-3 | CD02 | 44 | 20 | 10 | 1h 34m 56s | 2m 9.45s |
| part2 | 1A9X-3 | CD03 | 40 | 20 | 10 | 1h 37m 53s | 2m 26.82s |
| part2 | 1A9X | CD01 | 19 | 47 | 31 | 56m 12s | 2m 57.47s |
| part2 | 1A9X | CD02 | 44 | 47 | 31 | 3h 4m 19s | 4m 11.34s |
| part2 | 1A9X | CD03 | 40 | 47 | 31 | 1h 48m 27s | 2m 42.68s |
| part2 | 1B37 | CD01 | 16 | 32 | 10 | 29m 47s | 1m 51.69s |
| part2 | 1B37 | CD02 | 35 | 32 | 10 | 2h 16m 46s | 3m 54.46s |
| part2 | 1B37 | CD03 | 51 | 32 | 10 | 2h 42m 18s | 3m 10.94s |
| part2 | 1B37 | CD04 | 70 | 32 | 10 | 2h 52m 50s | 2m 28.14s |
| part2 | 1B37 | CD05 | 49 | 32 | 10 | 2h 14m 9s | 2m 44.27s |
| part2 | 1B37 | CD06 | 50 | 32 | 10 | 3h 19m 21s | 3m 59.22s |
| part2 | 1B37 | CD07 | 43 | 32 | 10 | 1h 57m 29s | 2m 43.93s |
| part2 | 1B37 | CD08 | 48 | 32 | 10 | 2h 37m 39s | 3m 17.06s |
| part2 | 1B37 | CD09 | 61 | 32 | 10 | 2h 42m 6s | 2m 39.44s |
| part2 | 1B37 | CD10 | 78 | 32 | 10 | 3h 9m 52s | 2m 26.05s |
| part2 | 1B37 | CD11 | 59 | 32 | 10 | 2h 45m 31s | 2m 48.32s |
| part2 | 1BRT | CD01 | 40 | 7 | 4 | 1h 15m 9s | 1m 52.72s |
| part2 | 1BRT | CD02 | 78 | 7 | 4 | 1h 37m 9s | 1m 14.73s |
| part2 | 1BRT | CD03 | 87 | 7 | 4 | 1h 43m 37s | 1m 11.46s |
| part2 | 1EA5 | CD01 | 36 | 26 | 10 | 1h 14m 9s | 2m 3.58s |
| part2 | 1EA5 | CD02 | 57 | 26 | 10 | 2h 14m 40s | 2m 21.75s |
| part2 | 1HNJ | CD01 | 20 | 89 | 51 | 53m 9s | 2m 39.45s |
| part2 | 1HNJ | CD02 | 40 | 89 | 51 | 39m 58s | 59.95s |
| part2 | 1HNJ | CD03 | 71 | 89 | 51 | 1h 2m 28s | 52.79s |
| part2 | 1I88-2 | CD01 | 19 | 91 | 54 | 49m 51s | 2m 37.42s |
| part2 | 1I88-2 | CD02 | 45 | 91 | 54 | 2h 13m 4s | 2m 57.42s |
| part2 | 1I88-2 | CD03 | 46 | 91 | 54 | 1h 39m 10s | 2m 9.35s |
| part2 | 1I88-2 | CD04 | 61 | 91 | 54 | 3h 19m 8s | 3m 15.87s |

| part | protein | tunnel | path length | ligand atoms | ligand heavy atoms | total time | avg time/frame |
|---|---|---|---|---|---|---|---|
| part2 | 1I88-2 | CD05 | 39 | 91 | 54 | 1h 45m 41s | 2m 42.59s |
| part2 | 1I88 | CD01 | 19 | 101 | 59 | 44m 48s | 2m 21.47s |
| part2 | 1I88 | CD02 | 45 | 101 | 59 | 1h 55m 53s | 2m 34.51s |
| part2 | 1I88 | CD03 | 46 | 101 | 59 | 1h 51m 46s | 2m 25.78s |
| part2 | 1I88 | CD04 | 61 | 101 | 59 | 2h 23m 4s | 2m 20.72s |
| part2 | 1I88 | CD05 | 39 | 101 | 59 | 2h 7m 37s | 3m 16.33s |
| part2 | 1M1N | CD01 | 24 | 47 | 31 | 1h 26m 54s | 3m 37.25s |
| part2 | 1M1N | CD02 | 55 | 47 | 31 | 4h 17m 50s | 4m 41.27s |
| part2 | 1M1N | CD03 | 55 | 47 | 31 | 3h 40m 7s | 4m 0.13s |
| part2 | 1PBE | CD01 | 32 | 15 | 10 | 1h 15m 15s | 2m 21.09s |
| part2 | 1PBE | CD02 | 25 | 15 | 10 | 1h 27s | 2m 25.08s |
| part2 | 1PBE | CD03 | 29 | 15 | 10 | 57m 56s | 1m 59.86s |
| part2 | 1PBE | CD04 | 58 | 15 | 10 | 2h 2m 31s | 2m 6.74s |
| part2 | 1PBE | CD05 | 48 | 15 | 10 | 1h 51m 11s | 2m 18.98s |
| part2 | 1VA4 | CD01 | 18 | 18 | 10 | 33m 58s | 1m 53.22s |
| part2 | 1VA4 | CD02 | 25 | 18 | 10 | 53m 29s | 2m 8.36s |
| part2 | 1VA4 | CD03 | 27 | 18 | 10 | 39m 30s | 1m 27.78s |
| part2 | 1YGE | CD01 | 50 | 51 | 20 | 3h 29m 6s | 4m 10.92s |
| part2 | 1YGE | CD02 | 58 | 51 | 20 | 3h 38m 31s | 3m 46.05s |
| part2 | 1YGE | CD03 | 51 | 51 | 20 | 3h 11m 21s | 3m 45.12s |
| part2 | 1YGE | CD04 | 76 | 51 | 20 | 5h 5m 40s | 4m 1.32s |
| part2 | 1YGE | CD05 | 88 | 51 | 20 | 5h 21m 38s | 3m 39.30s |
| part2 | 1YGE | CD06 | 92 | 51 | 20 | 7h 6m 37s | 4m 38.23s |
| part2 | 1YGE | CD07 | 160 | 51 | 20 | 10h 30m 42s | 3m 56.51s |
| part2 | 1YGE | CD08 | 210 | 51 | 20 | 11h 1m 31s | 3m 9.00s |
| part2 | 1YRC | CD01 | 24 | 27 | 11 | 43m 24s | 1m 48.50s |
| part2 | 1YRC | CD02 | 43 | 27 | 11 | 1h 21m 25s | 1m 53.60s |
| part2 | 1YRC | CD03 | 46 | 27 | 11 | 1h 17m 34s | 1m 41.17s |
| part2 | 2AYL | CD01 | 41 | 53 | 22 | 1h 31m 11s | 2m 13.44s |

| part | protein | tunnel | path length | ligand atoms | ligand heavy atoms | total time | avg time/frame |
|---|---|---|---|---|---|---|---|
| part2 | 2AYL | CD02 | 71 | 53 | 22 | 2h 35m 53s | 2m 11.73s |
| part2 | 2AYL | CD03 | 58 | 53 | 22 | 1h 57m 54s | 2m 1.97s |
| part2 | 2AYL | CD04 | 67 | 53 | 22 | 2h 42m 34s | 2m 25.58s |
| part2 | 2AYL | CD05 | 81 | 53 | 22 | 3h 12m 18s | 2m 22.44s |
| part2 | 2AYL | CD06 | 85 | 53 | 22 | 2h 59m 20s | 2m 6.59s |
| part2 | 2AYL | CD07 | 57 | 53 | 22 | 2h 54m 0s | 3m 3.16s |
| part2 | 2AYL | CD08 | 95 | 53 | 22 | 3h 55m 57s | 2m 29.02s |
| part2 | 2AYL | CD09 | 62 | 53 | 22 | 2h 47m 58s | 2m 42.55s |
| part2 | 2C3N | CD01 | 25 | 36 | 20 | 1h 20m 17s | 3m 12.68s |
| part2 | 2C3N | CD02 | 33 | 36 | 20 | 1h 27m 46s | 2m 39.58s |
| part2 | 2C3N | CD03 | 41 | 36 | 20 | 1h 40m 13s | 2m 26.66s |
| part2 | 2IID | CD01 | 34 | 23 | 12 | 1h 12m 0s | 2m 7.06s |
| part2 | 2IID | CD02 | 61 | 23 | 12 | 2h 52m 22s | 2m 49.54s |
| part2 | 2IID | CD03 | 55 | 23 | 12 | 2h 40m 38s | 2m 55.24s |
| part2 | 2IID | CD04 | 32 | 23 | 12 | 1h 39m 45s | 3m 7.03s |
| part2 | 2IID | CD05 | 32 | 23 | 12 | 1h 19m 4s | 2m 28.25s |
| part2 | 2IID | CD06 | 37 | 23 | 12 | 1h 53m 11s | 3m 3.54s |
| part2 | 2IID | CD07 | 30 | 23 | 12 | 1h 21m 1s | 2m 42.03s |
| part2 | 2IID | CD08 | 46 | 23 | 12 | 2h 11m 56s | 2m 52.09s |
| part2 | 2IID | CD09 | 57 | 23 | 12 | 2h 11m 8s | 2m 18.04s |
| part2 | 2IID | CD10 | 70 | 23 | 12 | 2h 7m 4s | 1m 48.91s |
| part2 | 2IID | CD11 | 68 | 23 | 12 | 2h 4m 32s | 1m 49.88s |
| part2 | 2IID | CD12 | 107 | 23 | 12 | 2h 35m 29s | 1m 27.19s |
| part2 | 2INC | CD01 | 47 | 15 | 7 | 1h 3m 32s | 1m 21.11s |
| part2 | 2INC | CD02 | 26 | 15 | 7 | 58m 48s | 2m 15.69s |
| part2 | 2INC | CD03 | 66 | 15 | 7 | 2h 36m 16s | 2m 22.06s |
| part2 | 2INC | CD04 | 82 | 15 | 7 | 2h 54m 42s | 2m 7.83s |
| part2 | 2Q9O-2 | CD01 | 35 | 58 | 32 | 4h 47m 43s | 8m 13.23s |
| part2 | 2Q9O-2 | CD02 | 50 | 58 | 32 | 4h 3m 2s | 4m 51.64s |

| part | protein | tunnel | path length | ligand atoms | ligand heavy atoms | total time | avg time/frame |
|---|---|---|---|---|---|---|---|
| part2 | 2Q9O-2 | CD03 | 42 | 58 | 32 | 4h 17m 2s | 6m 7.19s |
| part2 | 2Q9O-3 | CD01 | 35 | 17 | 9 | 1h 46m 40s | 3m 2.86s |
| part2 | 2Q9O-3 | CD02 | 50 | 17 | 9 | 2h 2m 59s | 2m 27.58s |
| part2 | 2Q9O-3 | CD03 | 42 | 17 | 9 | 1h 32m 30s | 2m 12.14s |
| part2 | 2Q9O-4 | CD01 | 35 | 46 | 26 | 1h 42m 26s | 2m 55.60s |
| part2 | 2Q9O-4 | CD02 | 50 | 46 | 26 | 3h 34m 28s | 4m 17.36s |
| part2 | 2Q9O-4 | CD03 | 42 | 46 | 26 | 2h 25m 29s | 3m 27.83s |
| part2 | 2Q9O | CD01 | 35 | 21 | 11 | 1h 35m 24s | 2m 43.54s |
| part2 | 2Q9O | CD02 | 50 | 21 | 11 | 2h 4m 28s | 2m 29.36s |
| part2 | 2Q9O | CD03 | 42 | 21 | 11 | 1h 53m 46s | 2m 42.52s |
| part2 | 2SQC | CD01 | 45 | 80 | 30 | 2h 16m 4s | 3m 1.42s |
| part2 | 2SQC | CD02 | 54 | 80 | 30 | 2h 27m 34s | 2m 43.96s |
| part2 | 2SQC | CD03 | 48 | 80 | 30 | 2h 26m 36s | 3m 3.25s |
| part2 | 2SQC | CD04 | 42 | 80 | 30 | 1h 50m 20s | 2m 37.62s |
| part2 | 2SQC | CD05 | 36 | 80 | 30 | 1h 50m 18s | 3m 3.83s |
| part2 | 2SQC | CD06 | 58 | 80 | 30 | 2h 58m 58s | 3m 5.14s |
| part2 | 2SQC | CD07 | 88 | 80 | 30 | 4h 22m 15s | 2m 58.81s |
| part2 | 3C6X | CD01 | 18 | 13 | 6 | 32m 36s | 1m 48.67s |
| part2 | 3TTV | CD01 | 24 | 4 | 2 | 1h 7m 42s | 2m 49.25s |
| part2 | 3TTV | CD02 | 30 | 4 | 2 | 1h 24m 31s | 2m 49.03s |
| part2 | 3TTV | CD03 | 66 | 4 | 2 | 2h 13m 5s | 2m 0.98s |
| part2 | 3TTV | CD04 | 68 | 4 | 2 | 2h 14m 51s | 1m 58.99s |
| part2 | 3TTV | CD05 | 74 | 4 | 2 | 2h 24m 57s | 1m 57.53s |
| part2 | 3TTV | CD06 | 85 | 4 | 2 | 2h 58m 26s | 2m 5.95s |

### Input JSON configuration reference

Both entry points read a single JSON config:

```
python lpf/lpf_coarse_path.py  run.json      # CoarsePathRunner
python lpf/lpf_trajectory.py       run.json      # TrajectoryRunner
```

Any value can be overridden on the command line with dot-notation:

```
python lpf/lpf_trajectory.py run.json --set parameters.step=0.5 --set rosetta_scripts.nstruct_main=20
```

The **Used by** column below is C (coarse path), T (trajectory), or C+T. See unused_config_settings.md for the keys each script *ignores*. examples/*/run.json are minimal working configs – one per run type.

---

#### Top level

| Key | Used by | Description |
|---|---|---|
| `rosetta` | C+T | Absolute path to the rosetta_scripts binary. If n_proc > 1 the filename must contain .mpi.. |
| `general_files` | C+T | Directory containing VRT1.params (the virtual-atom residue type used for all constraint anchor points). |
| `extra_files` | C+T | List of extra files copied into every Rosetta working directory, e.g. cofactor params (["../files/HEM.params"]). Use [] if none. |
| `input` | C+T | Input structures and paths – see below. |
| `parameters` | C+T | Algorithm and constraint parameters – see below. |
| `rosetta_scripts` | C+T | Rosetta protocol files and run sizes – see below. |
| `output` | C+T | Output location and run behaviour – see below. |

---

#### `input`

| Key | Used by | Description |
|---|---|---|
| `guide_path` | C+T | Path to the guide curve: either a whitespace-delimited x y z text file or a .pdb (ATOM/HETATM coordinates). **Coarse** uses only the first and last point; the route between them is generated by recursive bisection. **Trajectory** fits a smoothing spline through all points and resamples it at step spacing. |
| `ligand_files` | C+T | Directory holding lig.params (Rosetta params for the ligand; its NBR_ATOM is the atom tracked along the path) and lig_conformers.pdb (pre-generated conformer library). |

| Key | Used by | Description |
|---|---|---|
| `ligand_chain` | C+T | Chain ID of the ligand in the input PDBs (e.g. "X"). |
| `start_pdb` | C+T | Starting complex (protein + ligand in one file). Provide **either** this **or** the start_protein_pdb + start_ligand_pdb pair. |
| `start_protein_pdb` | C+T | Protein-only starting PDB. Merged with start_ligand_pdb into start_merged.pdb at run time. |
| `start_ligand_pdb` | C+T | Ligand-only starting pose PDB, in the frame of start_protein_pdb. |
| `header_pdb` | C+T | *Optional.* File whose lines are prepended to every generated task.pdb (membrane spans, symmetry records, etc.). Omit or set false if unused. |

---

### `parameters`

#### Path geometry

| Key | Used by | Description |
|---|---|---|
| `step` | C+T | Target spacing between points along the path, in Å. **Coarse:** bisection subdivides a segment until both half-gaps are < step. **Trajectory:** spline is resampled to windows step apart. |
| `smooth` | T | Smoothing factor s passed to scipy.interpolate.splprep for the guide-path spline. Larger = smoother / less faithful to the raw points. The trajectory run prints the max turning angle plus what 2 times and 5 times this value would give. |
| `endpoint_jitter` | C | If true, displace the path endpoint within a disc (radius endpoint_jitter_radius) perpendicular to the path axis, so independent runs land at slightly different endpoints – improves coverage when many runs are pooled and clustered. |
| `endpoint_jitter_radius` | C | Disc radius in Å for endpoint_jitter. Ignored unless endpoint_jitter is true. |

#### Conformer sampling

| Key | Used by | Description |
|---|---|---|
| `conformer_subsampling` | C+T | If true, before each Rosetta call filter lig_conformers.pdb down to conformers within conformer_rmsd_threshold of the current ligand pose (speeds up sampling for flexible ligands). If false, the full library is used every time. |
| `conformer_rmsd_threshold` | C+T | RMSD cutoff (Å) for conformer_subsampling. Ignored unless subsampling is on. |

#### RMSD (shape) constraint

Restrains each ligand atom toward its position in the incoming reference pose, via one `AtomPair` per atom to a per-atom ghost virtual atom. Acts as a soft whole-ligand RMSD restraint.

| Key | Used by | Description |
| --- | --- | --- |
| `rmsd_constraint` | C+T | Master on/off switch. (Always forced false for the dock-zero / window-0 dock step, where the incoming pose is not yet trustworthy.) |
| `rmsd_constraint_type` | C+T | "harmonic" (spring to the reference position) or "bounded" (flat well then quadratic). |
| `rmsd_constraint_bound` | C+T | Width (Å) of the flat region; used only when rmsd_constraint_type == "bounded". |
| `rmsd_constraint_sd` | C+T | Base standard deviation (Å); the effective per-atom weight is sd·√N. **Coarse** additionally scales it by √(dL/step) per node (looser over larger gaps). **Trajectory** retries a failed window with sd·0.2, sd·0.04. |
| `rmsd_constraint_hydrogens` | C+T | If false, hydrogens are excluded from the restraint. |

#### In-plane-deviation constraint (ipdc)

One `AtomPair` between the ligand `NBR_ATOM` and the target point for the current step, keeping the ligand near the guide point / inside the bisection plane.

| Key | Used by | Description |
| --- | --- | --- |
| `in_plane_dev_constraint` | C+T | Master on/off switch. |
| `ipdc_type` | C+T | "bounded" or "harmonic". **Coarse always uses "bounded"** – any other value is ignored (with a warning). |
| `ipdc_d0` | T | Flat-well radius (Å) for the bounded form. **Coarse ignores the config value** and recomputes it per node as 0.5·dL (dL = node distance to its target). |
| `ipdc_sd` | T | Constraint standard deviation (Å). **Coarse ignores the config value** and recomputes it per node as 0.1·dL. |
| `end_padding` | T | Distance (Å) over which the in-plane constraint is relaxed as the trajectory approaches the endpoint: the last int(end_padding/step) windows get a linearly shrinking ipdc_d0, and the terminal window snaps harmonically to the endpoint. 0 disables. **Coarse ignores this** (treated as 0, with a warning). |

#### Plane (coplanarity) constraint

Three `Dihedral` constraints tie the ligand `NBR_ATOM` into the plane of a virtual equilateral triangle placed in the bisection / guide plane.

| Key | Used by | Description |
| --- | --- | --- |
| `plane_constraint_point_extent` | C+T | Circumradius (Å) of the virtual triangle. Larger = stiffer, better-conditioned coplanarity restraint. |
| `plane_constraint_sd` | C+T | Standard deviation (radians) of the three dihedral constraints (target dihedral 0). Small values (~0.02) enforce the plane tightly. |

#### Direction (angle) constraints

Both target a straight angle (π); `bound` is the lower limit in radians (e.g. `2.65 rad ≈ 152°`, so the angle is held between 152° and 180°).

| Key | Used by | Description |
|---|---|---|
| `angle_constraint` | C+T | On/off. **Coarse:** Angle(prev_neighbour – NBR – next_neighbour) – keeps the ligand between its two path neighbours. **Trajectory:** Angle(prev_prev – prev – NBR) – penalises back-tracking. |
| `angle_constraint_type` | C+T | "bounded" or "harmonic". |
| `angle_constraint_bound` | C+T | Lower angle bound (radians); used only for "bounded". |
| `angle_constraint_sd` | C+T | Standard deviation (radians). |
| `next_angle_constraint` | C+T | On/off for the second angle term. **Coarse:** Angle(second-neighbour – neighbour – NBR) on each side. **Trajectory:** Angle(prev – NBR – next) – keeps forward progress aligned with the guide. |
| `next_angle_constraint_type` | C+T | "bounded" or "harmonic". |
| `next_angle_constraint_bound` | C+T | Lower angle bound (radians); "bounded" only. |
| `next_angle_constraint_sd` | C+T | Standard deviation (radians). |

#### Pose propagation & selection

| Key | Used by | Description |
|---|---|---|
| `memory_mode` | C+T | If true, the full best decoy (including any protein relaxation) is carried forward to the next step, so induced protein motion accumulates along the path. If false, only the ligand coordinates are transplanted onto the original protein via swap_lig_coords, and each step restarts from the same protein conformation. |
| `select_best_by` | C+T | Score used to pick the best decoy: "total_score" or "interface_delta_X". Constraint energies are subtracted before ranking. |
| `select_next_by` | T | Same choice, for the decoy that is *propagated* to the next window (may differ from the one reported). |
| `outlier_rmsd` | T | Reject decoys whose ligand NBR_ATOM has drifted more than this (Å) from the window's target position before selection. |
| `noise` | T | If true, add Gaussian noise (sd = step·0.33) to the projected previous point before each main window – helps escape local traps. |
| `skip` | T | If true, skip windows where the guide path folds back on itself (plane-crossing angle > 90°). |

#### Optional pre-docking

| Key | Used by | Description |
|---|---|---|
| `dock_zero` | C+T | If true, first dock the ligand onto the guide path's first point (using protocol_zero_dock / nstruct_zero) before the main algorithm starts. Use |

| Key | Used by | Description |
|---|---|---|
| | | when the starting structure has the ligand present but not yet positioned. Requires rosetta_scripts.protocol_zero_dock and rosetta_scripts.nstruct_zero. |

---

### `rosetta_scripts`

| Key | Used by | Description |
|---|---|---|
| `options` | C+T | Rosetta @-options (flags) file, copied into each working directory. |
| `protocol` | C | RosettaScripts XML run at every coarse-path node. |
| `protocol_zero` | T | XML for window 0 when dock_zero is false (typically a minimise-only protocol). |
| `protocol_main` | T | XML for all normal trajectory windows. |
| `protocol_stats` | T | XML run for the optional per-window statistics pass (see run_stats). |
| `protocol_zero_dock` | C+T | XML for the pre-docking step; required only when dock_zero is true. |
| `nstruct` | C | Decoys generated per coarse-path node. |
| `nstruct_main` | T | Decoys per trajectory window. |
| `nstruct_stats` | T | Decoys for the statistics pass; required only when run_stats is true. |
| `nstruct_zero` | C+T | Decoys for the pre-docking step; required only when dock_zero is true. |
| `n_proc` | C+T | MPI ranks. Must be a positive integer and ≤ the relevant nstruct. > 1 requires an .mpi. Rosetta binary. |
| `mpi_launcher` | C+T | Launcher command used when n_proc > 1 (e.g. "mpirun"). |

---

### `parameters.run_stats` / statistics pass

| Key | Used by | Description |
|---|---|---|
| `run_stats` | T | If true, after each window run protocol_stats (nstruct_stats decoys) on the selected pose and record separate stats score arrays into stats.pkl. |

---

### output

| Key | Used by | Description |
|---|---|---|
| path | C+T | Output directory. Created if absent; all per-node / per-window subdirectories and the final trajectory files are written here. |
| verbose | C+T | Extra progress logging. Default true. |
| archive | C+T | If true, zip -rm each node/window directory after it finishes, to save inodes. |
| checkpoint_interval | T | Save a resumable checkpoint every N completed windows. 0 disables. A run auto-resumes from checkpoint.pkl in path unless started with --fresh. |
| use_tmp | T | If true, execute each window in a temporary directory (RAM disc by default) and copy results back to path on completion. |
| tmp_path | T | Explicit temporary location for use_tmp. Empty string → try /dev/shm, then /run/shm. |

---

### Outputs written to `output.path`

**Coarse path:** coarsepath.txt (point sequence), trajectory.pdb, lig_trajectory.pdb, full_trajectory.pdb, and one numbered directory per node.

**Trajectory:** guide_path / spline / normals / input_path, nbr_trajectory.txt, total_scores.txt, interface_scores.txt, stats.pkl, lig_trajectory.pdb, full_trajectory.pdb, checkpoint.pkl, and one window_XXXX directory per window.

### Clustering of sampled paths

Pairwise distances between paths were computed as the frame-by-frame coordinate RMSD of the ligand, summed over frames with linearly decreasing weights that therefore put more weight into conformational differences at early frames, when the ligand is within the protein tunnel. If one path was shorter, each missing frame was assigned a fixed penalty of 0.5 Å. The negative of the resulting RMSD matrix was passed as a precomputed affinity matrix to affinity propagation (scikit-learn, default parameters). The interaction energy profile of each path (interface score per frame, 0.33 Å per frame) was then averaged within each cluster. Clusters of 10 or fewer paths were discarded. The remaining cluster averages were plotted against progression along the path (Å). The most populated cluster is drawn as a bold line, and the other retained clusters are drawn as thin, semi-transparent lines.

A script for clustering is shared on zenodo within 10.5281/zenodo.23183178.

### Protocol capture

This tutorial walks through a complete Ligand PathFinder (LPF) run – coarse path finding followed by a trajectory – on one worked example. Every input file used below is pre-made and shipped in this demo/ folder shared on zenodo as 10.5281/zenodo.23183178:

```
demo/
├── tutorial_1.md                ← this file
├── figures/                     ← figures referenced in the text
├── coarse_path/                 ← stage 2 (see §2)
│   ├── files/                   coarsepath.xml, options, VRT1.params
│   ├── ligand/                  lig.params, lig_conformers.pdb
│   ├── path                     two-point guess direction
│   ├── header.pdb               empty (no metal constraints)
│   ├── start.pdb                minimized BChE-POX complex
│   ├── run.json                 run configuration
│   └── run/                     the finished coarse-path run
└── trajectory/                  ← stage 3 (see §3)
    ├── files/                   main_standard.xml, zero_minimize.xml,
stats.xml, options, VRT1.params
    ├── ligand/                  lig.params, lig_conformers.pdb
    ├── path                     the coarse path from stage 2 (coarsepath.txt)
    ├── header.pdb               empty
    ├── start.pdb                same minimized complex
    ├── run.json                 run configuration
    └── run/                     the finished trajectory run
```

Please note any parts that could benefit from additional automation and send feature requests to the developers.

#### Requirements

- A Conda environment built from environment.yml at the repository root (conda env create -f environment.yml && conda activate ligpf). The Python side only needs numpy and scipy; zip is used when output.archive is on.
- A Rosetta rosetta_scripts binary – its path goes into the "rosetta" key of run.json. An MPI launcher (e.g. mpirun) is only needed if you set rosetta_scripts.n_proc > 1 (and then the binary must be an .mpi. build).

#### 1. System preparation

For this tutorial we use **human butyrylcholinesterase (BChE)**. It has a deep binding gorge that is well suited to exploration with Ligand PathFinder. As a ligand we use paraoxon (POX), a covalent inhibitor of the enzyme that uses its catalytic machinery for the first step of the acylation reaction.

Ligand PathFinder starts from a PDB file of the enzyme–ligand complex in a functionally relevant conformation / binding pose:

- For enzyme-substrate complexes (a covalent inhibitor can be treated as a substrate for a partial reaction) – the pre-reactive state.

- If the question concerns product release – the product placed in the immediate after-reaction pose.
- For enzyme-inhibitor complexes – the functional binding pose of the inhibitor.
- For complex cases (multiple substrates, cofactors, binding-triggered conformational changes) – understand the sequence of events from fundamental studies (what binds first, to which protein conformation, what unbinds first, how the conformation changes) and reduce the task to the elementary step.

#### 1.1 Binding pose

Constructing the protein-ligand(s) complex is the first prerequisite for Ligand PathFinder, and it is **not** solved within it. Any docking tool or approach can be used at this stage: AutoDock Vina, Rosetta, GLIDE, DiffDock, AF3 / chai-1 / boltz-1 prediction, manual construction. Manual verification of the adequacy of the result is recommended.

Here we use a chai-1 prediction, since it agrees perfectly with our previous results:

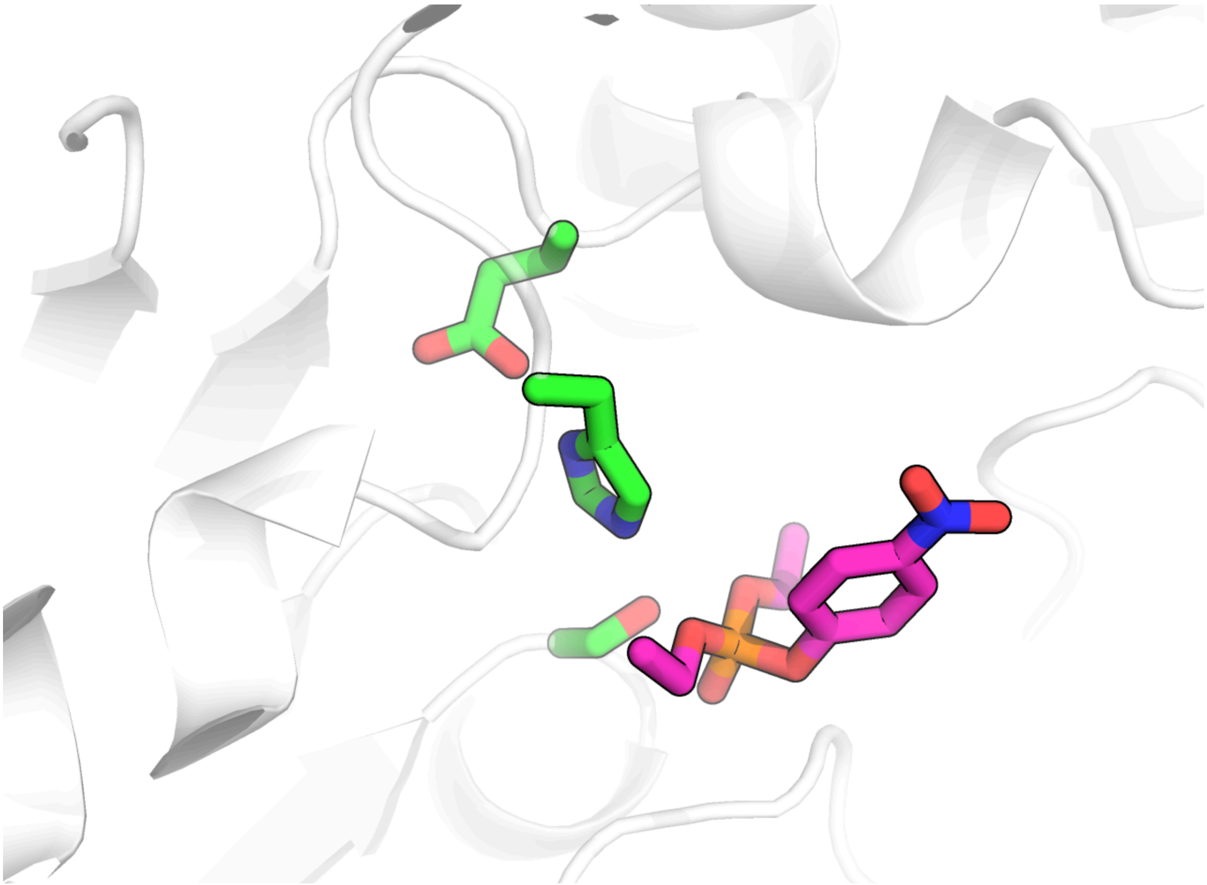

#### 1.2 Rosetta parameters for the ligand

**1.2.1.** First, we need a standalone **sdf** file for the ligand with protons and correctly specified bond orders. For some docking tools this file already exists. We can work with the ligand we have from the docking stage, or use it only as guidance and create a standalone protonated file independently.

If we work with a chai-generated cif file:

- We can protonate in PyMOL, then double-check protonation and bond orders and correct them in the builder tool.
- We can use PyMOL to save only the ligand in a standalone file and use Avogadro to manually add protons and correct bond orders.
- Or use any proton-adding tool on this file (SPORES, OpenEye, OpenBabel).

If we create the file anew:

- Build the molecule in Avogadro and then minimize.
- Download the pre-built sdf from a database.
- Use any SMILES-to-sdf converter tool with subsequent energy minimization.

For POX, an sdf file was available on PubChem. It is Kekulized. For the Rosetta parameter creation steps it is better to specify aromaticity explicitly – i.e. change bond types from alternating 1, 2 to 4 for the whole aromatic ring. We do this manually.

**1.2.2.** We need a library of conformers for the ligand in **sdf** format. A number of tools can be used. The default RosettaLigand way is to use BCL as described in the tutorials (here, step 3.2). Other tools are RDKit, CREST, OpenEye. Here we used BCL:

```
bcl molecule:ConformerGenerator -ensemble_filenames pox.sdf \
    -conformers_single_file pox_conf.sdf -explicit_aromaticity
```

**1.2.3.** We create Rosetta parameters for the ligand in the standardized way:

```
rosetta/main/source/scripts/python/public/molfile_to_params.py \
    -n lig -p lig --conformers-in-one-file pox_conf.sdf
```

We get several crucial files:

- lig.pdb
- lig.params
- lig_conformers.pdb

lig.params and lig_conformers.pdb are needed directly for Ligand PathFinder runs. lig.pdb is needed to modify the complex PDB file, since we need a starting PDB with the ligand not only in the correct pose but also with protons and all atoms **in the correct order**.

**1.2.4.** We create a starting protein-ligand complex PDB that works with the Rosetta parameter files. One way is PyMOL:

```
set retain_order, 1
```

Then open the protein-ligand docking result and lig.pdb alongside. Manually adjust the position and conformation of lig from lig.pdb to match the pose from the previous docking (chai prediction). Finally, output the protein and the ligand from lig.pdb into a new PDB file, complex.pdb.

> In demo/*/ligand/ you will find the resulting lig.params and lig_conformers.pdb for POX. The ligand's NBR_ATOM (declared in lig.params) is O1 – this is the atom LPF tracks along the path. Note that LPF now reads the neighbour atom from lig.params directly; there is no longer a ligand_nbr key in run.json.

#### 1.3 Starting file for Ligand PathFinder

We adjust the atomic model to the Rosetta force field with a relaxation or a minimization:

```
rosetta/main/source/bin/minimize.static.linuxgccrelease -s complex.pdb \
    -extra_res_fa lig.params -restore_pre_talaris_2013_behavior
```

Manual inspection of the relaxed structure is highly recommended. It is also recommended to run several attempts (-nstruct <N>) and pick the best for the subsequent LPF run.

> **On generating the starting pose automatically.** LPF now has a dock_zero option: when it is set, before the main routine starts, the ligand is docked onto the first point of the guide path (using rosetta_scripts.protocol_zero_dock / nstruct_zero). This is useful when the starting structure contains the ligand but not yet in position. **We still strongly recommend preparing and validating the binding pose separately** – it is far too important a step to delegate to any algorithm alone. In this tutorial we keep dock_zero: false and provide a hand-prepared, minimized start.pdb. When dock_zero is off, window 0 of the trajectory is still passed through protocol_zero (zero_minimize.xml), which only settles the supplied pose into the force field.

*Optional:* relaxation of an enzyme-substrate complex in Rosetta is a fine and peaky process. It may not be solved automatically for a particular system. One option is to first relax the protein without the ligand, then superpose it back, transfer the ligand, and repeat the relaxation/minimization. Another is to use custom relaxation scripts (particularly for metalloenzymes) through the XML RosettaScripts interface, potentially with a Cartesian score function and manually curated restraints, to reliably model pre-reactive poses.

*Optional:* enzymes sometimes have non-standard protonation patterns and crucial water molecules. These cases require manual attention. Manually specifying a His tautomer, Asp/Glu/His protonation state, or neutral termini is currently not possible in Rosetta; please contact the core developers if your systems require it.

In the end we copy the selected best configuration as start.pdb. **It is crucial that in this file the ligand is placed after the protein** – see demo/coarse_path/start.pdb.

### 2. Coarse path finding

The main Ligand PathFinder routine needs a **guess path**. It can be created manually or with third-party software such as CAVER or AQUA-DUCT. In that case the path is a series of points in 3D space written line by line in a text file, starting from the coordinates of the ligand's NBR_ATOM:

```
1.104 0.834 0.452
1.054 0.800 0.552
1.000 0.777 0.587
…
0.245 0.345 0.923
```

If you already have such a path, skip to §3 and use it as the trajectory path file.

Ligand PathFinder also has a routine to create guess paths by itself – this is the **coarse path** stage, and it is what we use here.

#### 2.1 Preparation

The folder structure for a coarse-path run (demo/coarse_path/):

```
coarse_path/
├── files/
│   ├── VRT1.params
│   ├── coarsepath.xml
│   └── options
├── ligand/
│   ├── lig.params
│   └── lig_conformers.pdb
├── path
├── header.pdb
├── start.pdb
└── run.json
```

files/ holds *general* LPF files that do not depend on the system. The XML instructions and the options file are generic; change them only if you know exactly what you want to happen differently. In options, the packing/scoring flags are standard RosettaLigand settings (the per-calculation sampling depth is controlled by rosetta_scripts.nstruct in run.json, not here).

> files/ **does not have to be unique per project.** These are the same files shipped with the repository in rosettaXML_files/. You can copy them into each run folder (as here), or simply point general_files and the rosetta_scripts.* paths in run.json straight at rosettaXML_files/. The coarse protocol XML lives in demo/coarse_path/files/coarsepath.xml.

**Non-standard residues.** If your system needs extra parameters for a residue you built yourself (non-canonical amino acid such as carboxy-lysine, a non-standard cofactor), place the params file in files/, add its name to the options file (-extra_res_fa), and add its path to extra_files in run.json.

ligand/ holds the system-specific ligand files from §1.2.

path and run.json are also system-specific. For the coarse stage, path has exactly two lines. The first point is the location of the ligand's NBR_ATOM; the second roughly defines the direction of unbinding. The endpoint is jittered internally (endpoint_jitter), so repeated coarse-path runs with the same inputs yield a *collection* of guess paths that can be clustered, with the medoid taken forward to the trajectory stage.

For our BChE–POX case the two points are (shown as spheres):

```
5.755 -5.775 -3.752
9.156 -25.469 -1.308
```

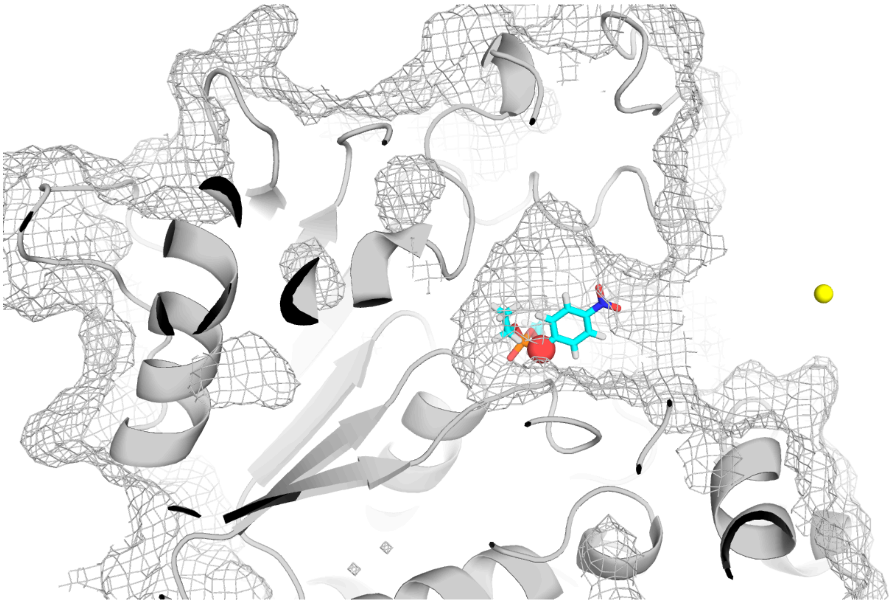

run.json holds all run parameters and the paths to run-related files. The same file structure is used for both the coarse-path and the trajectory scripts; see [docs/config_reference.md](docs/config_reference.md) for every key. demo/coarse_path/run.json:

```
{
  "rosetta": "<path_to>/rosetta_scripts.linuxgccrelease",
  "general_files": "../files",
  "extra_files": [],
  "input": {
    "guide_path": "../path",
    "ligand_files": "../ligand",
    "ligand_chain": "X",
    "start_pdb": "../start.pdb",
    "header_pdb": "../header.pdb"
  },
  "parameters": {
    "step": 2.0,
    "conformer_subsampling": false,
    "rmsd_constraint": true,
    "rmsd_constraint_type": "harmonic",
    "rmsd_constraint_sd": 1.5,
    "rmsd_constraint_hydrogens": false,
```

```
    "in_plane_dev_constraint": true,
    "ipdc_type": "bounded",
    "plane_constraint_sd": 0.02,
    "plane_constraint_point_extent": 7,
    "angle_constraint": true,
    "angle_constraint_type": "bounded",
    "angle_constraint_bound": 2.25,
    "angle_constraint_sd": 0.50,
    "next_angle_constraint": true,
    "next_angle_constraint_type": "bounded",
    "next_angle_constraint_bound": 1.57,
    "next_angle_constraint_sd": 0.25,
    "memory_mode": false,
    "select_best_by": "total_score",
    "endpoint_jitter": true,
    "endpoint_jitter_radius": 5.0,
    "dock_zero": false
  },
  "rosetta_scripts": {
    "options": "../files/options",
    "protocol": "../files/coarsepath.xml",
    "nstruct": 10,
    "n_proc": 1,
    "mpi_launcher": "mpirun"
  },
  "output": {
    "path": "./",
    "archive": true,
    "verbose": true
  }
}
```

It is recommended to leave everything in parameters as given here for all coarse-path runs.

header.pdb**.** Used only for metalloenzymes, to define metal-binding constraints that then will be prepended to all the starting pdb files for all tasks. BChE is metal-free, so we use an empty file (touch header.pdb) – demo/coarse_path/header.pdb.

#### 2.2 Run

Create a folder for the run, go there, and execute the script:

```
mkdir -p demo/coarse_path/run && cd demo/coarse_path/run
python <path_to_lpf>/lpf/lpf_coarse_path.py ../run.json
```

You start to get messages like:

```
STARTED: point 0
FINISHED: point 0
in 0:00:29.

STARTED: point 1
FINISHED: point 1
in 0:00:31.
```

which means the script is running correctly. The bisection tree subdivides the segment between the two endpoints until neighbouring points are closer than step (2.0 Å here), so the number of points depends on the path length. For this case it produced 15 nodes (17 points including the two endpoints) in a few minutes with nstruct: 10 on one core.

When it finishes you get:

- coarsepath.txt – xyz coordinates of the guess path. This is the input path for the next step (trajectory).
- full_trajectory.pdb – protein + ligand along the coarse path.
- lig_trajectory.pdb – ligand only.
- trajectory.pdb – the guess path as PDB points.

For the coarse stage the ligand/protein trajectories are not very useful on their own – the purpose here is only to produce a guess path. They are handy for debugging. The finished run is in demo/coarse_path/run/; the individual node directories are zipped (archive: true).

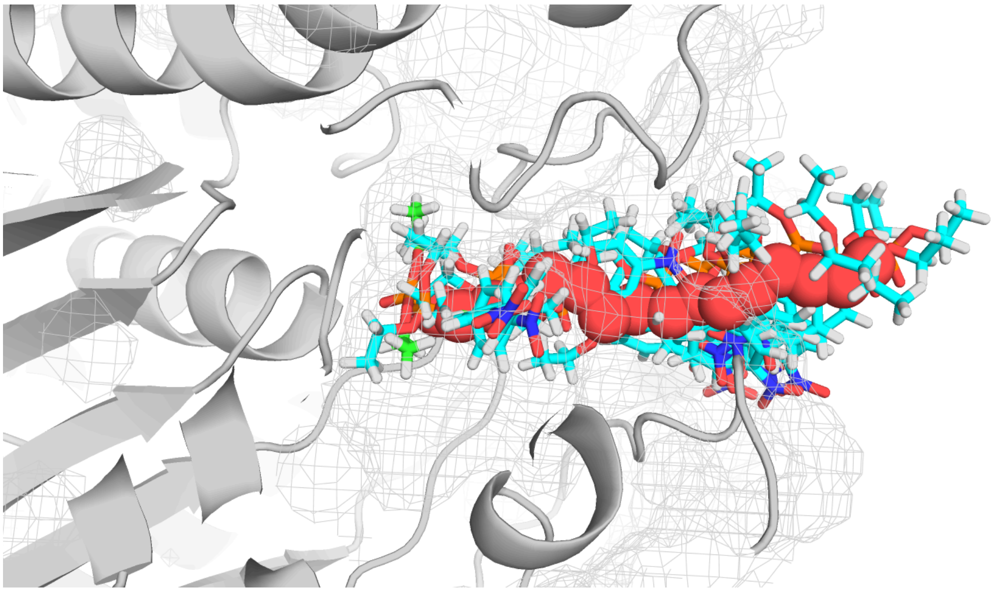

---

### 3. Trajectory

Create the same folder structure as for the coarse path (demo/trajectory/). You can reuse files/ and ligand/. The path file is now the *full* guess path, so we copy coarsepath.txt from §2 into demo/trajectory/path.

The trajectory needs a few extra generic XMLs in files/:

- main_standard.xml – the per-window protocol (was protocol.xml).
- zero_minimize.xml – window-0 protocol (protocol_zero); settles the start pose into the force field.
- stats.xml – optional second scoring pass (protocol_stats), only used if

run_stats: true.

Adjust run.json for the trajectory regime – demo/trajectory/run.json (only the parts that differ from §2.1 are highlighted in the comments):

```
{
  "rosetta": "<path_to>/rosetta_scripts.linuxgccrelease",
  "general_files": "../files",
  "extra_files": [],
  "input": {
    "guide_path": "../path",
    "ligand_files": "../ligand",
    "ligand_chain": "X",
    "start_pdb": "../start.pdb",
    "header_pdb": "../header.pdb"
  },
  "parameters": {
    "step": 0.5,
    "smooth": 0.5,
    "conformer_subsampling": true,
    "conformer_rmsd_threshold": 1.5,
    "rmsd_constraint": true,
    "rmsd_constraint_type": "harmonic",
    "rmsd_constraint_sd": 0.75,
    "rmsd_constraint_hydrogens": true,
    "in_plane_dev_constraint": true,
    "ipdc_type": "bounded",
    "ipdc_d0": 3.0,
    "ipdc_sd": 0.5,
    "end_padding": 5,
    "plane_constraint_sd": 0.02,
    "plane_constraint_point_extent": 5,
    "angle_constraint": true,
    "angle_constraint_type": "bounded",
    "angle_constraint_bound": 2.00,
    "angle_constraint_sd": 0.50,
    "next_angle_constraint": true,
    "next_angle_constraint_type": "bounded",
    "next_angle_constraint_bound": 1.57,
    "next_angle_constraint_sd": 0.25,
    "memory_mode": true,
    "run_stats": false,
    "select_best_by": "total_score",
    "select_next_by": "total_score",
    "noise": false,
    "outlier_rmsd": 2.0,
    "skip": true,
    "dock_zero": false
  },
  "rosetta_scripts": {
    "options": "../files/options",
    "protocol_zero": "../files/zero_minimize.xml",
    "protocol_main": "../files/main_standard.xml",
    "protocol_stats": "../files/stats.xml",
    "nstruct_main": 10,
    "nstruct_stats": 10,
    "n_proc": 1,
    "mpi_launcher": "mpirun"
  },
  "output": {
```

```
    "path": "./",
    "archive": true,
    "verbose": true,
    "checkpoint_interval": 10
  }
}
```

What changes relative to the coarse stage:

- step is small (0.5 Å)
- smooth is now active: the guess path is fitted with a smoothing spline (larger = smoother) and resampled at step spacing. The script prints the maximum turning angle and what 2× / 5× smoothing would give.
- conformer_subsampling is on, so before each window the conformer library is filtered to conformers within conformer_rmsd_threshold of the current pose.
- memory_mode: true – the full relaxed pose (including protein motion) is carried to the next window, so induced-fit changes accumulate along the path.
- ipdc_d0 / ipdc_sd / end_padding **are** honoured here (unlike the coarse stage). end_padding relaxes the in-plane constraint over the last few windows so the ligand can settle into bulk solvent.
- skip: true skips windows where the guide path folds back on itself.
- run_stats: false here to keep the run short; set it to true (with stats.xml) for a second, independent binding-energy estimate per window.
- checkpoint_interval: 10 writes a resumable checkpoint every 10 windows; rerun the same command to resume, or add --fresh to start over.

Create a working folder, go there, and execute:

```
mkdir -p demo/trajectory/run && cd demo/trajectory/run
python <path_to_lpf>/lpf/lpf_trajectory.py ../run.json
```

At start-up the script reports the guide-path curvature and suggests smoother alternatives – for this case:

```
Guide path curvature: max turn = 47.3° (smooth=0.5)
  informational: 2x smooth (1.0) → 29.2°,  5x smooth (2.5) → 15.2°
  WARNING: 1 step(s) exceed 25°; the angle constraint may not enforce
direction at these points:
    gp[25]: 47.3°
```

A sharp kink like this is where the ligand may struggle to follow the guide; if a run stalls or the ligand cuts the corner, raise smooth or run more coarse-path replicates and pick a straighter medoid.

Progress messages look like:

```
STARTED:  window 1/49
  cst:  angle=0.00  pair=1.19  dihedral=0.00  total=1.19
FINISHED: window 1/49
in 0:00:37.
d_to_plane 0.000
d_in_plane 0.245
```

d_to_plane / d_in_plane are how far the ligand's NBR_ATOM ended up from the guide plane and from the guide point – small values mean the constraints are holding. Window 0 is the fast zero_minimize.xml pass; the main windows here take ~30–40 s each with nstruct_main: 10 on one core (~30 min for the whole path). Windows where the guide path folds back on itself are skipped (skip: true):

```
SKIPPED:  window 37/49  (guide path turns back; plane angle = 167.1°)
SKIPPED:  window 38/49  (guide path turns back; plane angle = 173.5°)
```

For this case the spline gave 50 windows (0–49); 2 were skipped, leaving 48 scored windows. Finished windows are zipped to save space. The main analysis files (demo/trajectory/run/):

- interface_scores.txt – Rosetta binding-energy estimate (interface_delta_X) for each scored window. **Main file for analysis.**
- full_trajectory.pdb – unbinding trajectory, protein + ligand.
- lig_trajectory.pdb – ligand only.
- total_scores.txt – total complex energy per window (mostly for debug).
- nbr_trajectory.txt – NBR_ATOM xyz per scored window (debug).
- guide_path, spline, input_path, normals – path construction intermediates (debug).
- stats.pkl – per-window stats scores; populated only when run_stats: true.
- checkpoint.pkl – resume state (checkpoint_interval).

Plot the binding energy along the path with matplotlib, and inspect the per-window configurations in PyMOL. The profile below was made from this demo's own demo/trajectory/run/ output – interface_scores.txt against the cumulative distance travelled by the ligand NBR_ATOM (from nbr_trajectory.txt):

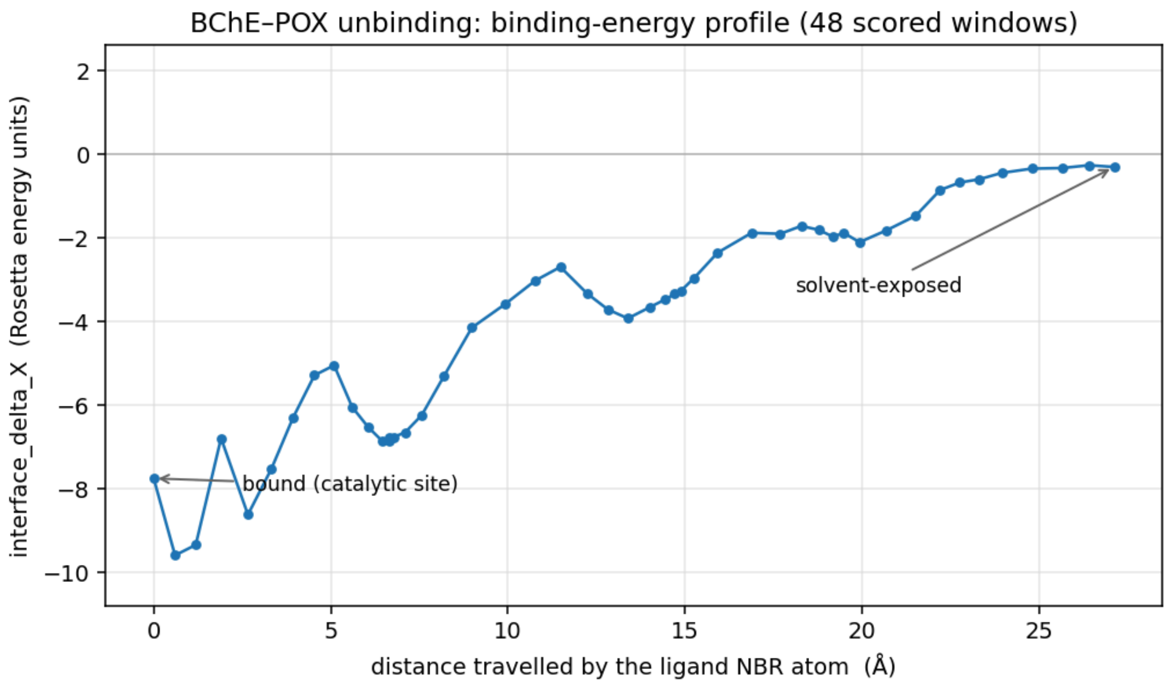

The bound state at the catalytic site is favourable (≈ −8 REU); the score climbs to ≈ 0 as POX leaves the ~27 Å gorge, i.e. this path releases on the order of ~7 REU of interface energy, with shallow local minima where the ligand pauses against gorge-wall residues. A single replicate is noisy – for quantitative work, pool interface_scores.txt over many replicates (§4).

> **Reproducing this.** The demo/coarse_path/run/ and demo/trajectory/run/ folders are the actual outputs of the two run.json files above against the current code (nstruct 10, one core: coarse ~5 min, trajectory ~35 min). Regenerate the figure after your own run with a two-line script (np.loadtxt('interface_scores.txt')).

---

### 4. (Optional) Replicates

This is a workflow note rather than part of the tutorial. Keeping one folder per run, with the paths in run.json pointing one level up, makes it easy to sample on a cluster through an sbatch array.

A minimal sbatch script (run.bash):

```
#!/bin/bash
#SBATCH -n 1
#SBATCH --cpus-per-task=1
#SBATCH --mem=5G
#SBATCH -t 720
#SBATCH -p paul,polaris
RUN_SCRIPT='<path_to>/python <path_to>/lpf/lpf_trajectory.py'
RUNDIR=repl-$SLURM_ARRAY_TASK_ID
mkdir $RUNDIR
cp run.json $RUNDIR
cd $RUNDIR
$RUN_SCRIPT run.json
```

Request as many replicas as you want:

```
sbatch -J task_name -a 0-29 run.bash
```

0-29 is the task range; each value is passed in as $SLURM_ARRAY_TASK_ID. This runs 30 independent replicas, each creating its own repl-* folder. Pool the per-window interface_scores.txt across replicas for a converged profile.

---

### 5. Adjustments

Running a single replica to completion first helps inform the set-up. Analysing the trajectory, you may notice:

1. The ligand is too stiff, it is mostly moving as a rigid body, not enough conformer sampling

    a. Check your initial conformer file, does it have enough different conformations? Are strained conformations (that might be nevertheless stabilized by protein) present there? Have you filtered by energy, or clustered by RMSD?
    b. Increase `conformer_rmsd_threshold`
    c. Increase `rmsd_constraint_sd`
    d. Consider turning off `rmsd_constraint_hydrogens`
    e. Decrease `angle_constraint_bound`
2. The ligand changes its conformation too sharply between steps
    a. Check suggestions from point 1, but in reverse
    b. Decrease the `step`
3. The protein is not flexible enough
    a. Check that you have `"memory_mode": true`
    b. Within the **"protocol.xml"**, increase `Calpha_restraints`
    c. Within the **"protocol.xml"**, increase cutoffs for `LigandAreas`
4. The protein assumes weird conformations
    a. Check suggestions from point 3, but in reverse
